# Structure and energy transfer of a minimal PSI–LHCI supercomplex with FNR binding from a terrestrial eustigmatophyte

**DOI:** 10.64898/2026.08.10.743057

**Authors:** Yi Luo, Kang Li, Quan Wen, Xiao-Meng Sun, Fang Zhao, Xin-Xiao Qu, Hao-Jie Wang, Jun Gao, Luo-Dong Huang, Yu-Zhong Zhang, Lu-Ning Liu, Long-Sheng Zhao

**Author notes:** Corresponding author: Long-Sheng Zhao, **Email:** (LSZ). Yi Luo, Kang Li, Quan Wen, and Yu-Zhong Zhang contributed equally to this work.

## Abstract

Soil microalgae endure harsh terrestrial stressors, such as intense light. Eustigmatophytes are an independent evolutionary branch within stramenopiles and occupy diverse aquatic and terrestrial environments, but the structural organization of their photosynthetic apparatus remains poorly understood. Here, we determined the cryo-electron microscopy structure of a photosystem I–light-harvesting complex I (PSI–LHCI) supercomplex bound with ferredoxin-NADP^+^ oxidoreductase (FNR) from the terrestrial eustigmatophyte *Vischeria stellata* at 2.44 Å resolution. The supercomplex contains a monomeric PSI core associated with only three LHCI subunits, representing the smallest PSI–LHCI reported among structurally characterized red-lineage PSI complexes composed of violaxanthin-Chl *a* proteins (VCPs). The three VCPIs with distinct structure features and arrangements form a compact belt along the PsaL–PsaI–PsaM side of PSI. The structure also resolves a 43-residue N-terminal segment of FNR (FNR-N) bound to the PSI stromal surface, which is stabilized by both a eustigmatophyte-conserved insertion in PsaL and the N-terminal region of PsaD. In contrast, the catalytic region of FNR was not resolved, suggesting conformational flexibility. Computational simulations indicate potential excitation-energy-transfer pathways connecting the three VCPI subunits to the PSI core and highlight lineage-specific pigments that maintain energetic connectivity within the exceptionally compact antenna. Our analysis further reveals conservation of FNR tethering despite pronounced diversification of antenna size and organization. These findings uncover a modular evolutionary principle in which PSI acceptor-side organization is retained while the light-harvesting antenna is extensively remodeled, providing a framework for understanding the diversification of photosynthetic energy conversion across ecological transitions.

## Introduction

Photosynthesis is one of the most important energy-conversion processes on Earth, and photosystem I (PSI) is a core component of the photosynthetic electron transport chain. In linear electron flow (LEF), PSI transfers electrons to ferredoxin (Fd), and ferredoxin-NADP^+^ oxidoreductase (FNR) accepts electrons from Fd and subsequently transfers them to NADP^+^ to produce NADPH, providing reducing power for carbon fixation (1). The topology of FNR is highly conserved: it consists of two independent domains connected by a linker loop, with the N-terminal domain responsible for binding the flavin adenine dinucleotide (FAD) cofactor and the C-terminal domain responsible for binding NADP^+^. Fd binds in a large groove between the two FNR domains (2).

In photosynthetic eukaryotes, transmembrane light-harvesting complex I (LHCI) proteins associate with the PSI core to form PSI–LHCI supercomplexes. Structures from red-lineage algae reveal exceptional diversity in antenna composition and organization, ranging from compact single-layer arrangements in red algae to large, multilayered assemblies in cryptophytes, haptophytes, diatoms, xanthophyte, brown algae, and dinoflagellates (3–21). Ochrophyte PSI antennae include RedCAP, Lhcr, Lhcf, and Lhcq-family proteins whose number and positions vary among lineages and growth conditions. This structural diversity provides a framework for evaluating how conserved PSI cores accommodate lineage-specific LHC antennae and highlights the divergent evolutionary adaptation strategies through which algal groups adapt photosynthetic energy conversion to distinct ecological niches.

Terrestrial algae are widely distributed in surface soils of forests, grasslands, deserts, and other terrestrial ecosystems (22). Compared with aquatic algae, terrestrial algae face stronger light, larger temperature fluctuations, and desiccating air stress during photosynthetic energy conversion (23). Structural and functional analyses of their photosystems are therefore critical for investigating the strategies by which their photosynthetic energy conversion adapts to the specialized light environment of soil habitats. However, high-resolution structures of PSI–LHCI supercomplexes from terrestrial algae remain scarce. To date, the only reported structure of soil algal PSI–LHCI is that of *Chlorella ohadii*, a green alga isolated from desert soil crusts (24). Additional structural studies are needed to clarify how the photosynthetic machinery of soil algae functions in terrestrial environments.

Eustigmatophyceae constitute a distinct lineage within Ochrophyta, the photosynthetic branch of stramenopiles (25), and are placed as the sister group to the Raphidophyceae–Phaeophyceae–Xanthophyceae (RPX) clade (26). Eustigmatophytes are globally distributed and occur predominantly in terrestrial and freshwater environments, whereas members of *Nannochloropsis* and *Microchloropsis* are found mainly in marine or brackish environments (27). The pigment composition of eustigmatophytes shows distinct lineage-specific features (28): eustigmatophytes generally lack chlorophyll (Chl) *c* and adopt Chl *a* as their major Chls, and their carotenoid composition is dominated by violaxanthin (29). Accordingly, their light-harvesting antennae are primarily composed of violaxanthin-Chl *a* proteins (VCPs) (30).

The recently reported structure of the marine eustigmatophyte *Nannochloropsis oceanica* revealed a PSI complex with nine VCPIs and an FNR N-terminal extension docked to the stromal PSI surface (31). Whether this FNR-docking architecture is restricted to *Nannochloropsis* or conserved across Eustigmatophyceae, and how PSI antenna organization differs in terrestrial representatives, remained unclear. Here, we determined the 2.44 Å cryo-electron microscopy (cryo-EM) structure of PSI–VCPI from the terrestrial eustigmatophyte *Vischeria stellata* (32). The complex contains only three peripheral antennas, and an FNR N-terminal segment was identified to bind to the stromal PSI surface. Comparative structural and sequence analyses show conserved determinants of FNR docking and extensive divergence in antenna organization between terrestrial and marine eustigmatophytes. Structure-based computational simulations further indicate candidate excitation-energy-transfer (EET) pathways within the minimal PSI–LHCI supercomplex. These findings shed light on the adaptive strategies supporting photosynthetic energy conversion and dissipation of soil microalgae under terrestrial environment.

## Results and Discussion

### Overall architecture

The PSI–VCPI supercomplex purified from *V. stellata* was characterized by absorption spectroscopy, SDS-PAGE, mass spectrometry, and pigment analysis (Fig. S1). The structure was determined by single-particle cryo-EM at an overall resolution of 2.44 Å (Fig. S2, Table S1). VsPSI–VCPI represents a minimal PSI–LHCI supercomplex compared to those of other red-lineage PSI–LHCI complexes, containing a monomeric PSI core and three peripheral VCPIs, termed VCPI-1, VCPI-2, and VCPI-3 (Fig. 1). The three VCPIs form a continuous antenna belt on the PsaL–PsaI–PsaM side of the PSI core. This antenna system is substantially smaller than those of recently reported red-lineage PSI complexes, including the nine VCPI antennae from *N. oceanica* (8, 13, 14, 18, 31). On the stromal side, a 43-residue N-terminal segment of FNR is resolved near PsaB, PsaD, and PsaL (Fig. 1). In total, the PSI–VCPI supercomplex contains 114 Chl *a* molecules, 15 *β*-carotenes, 9 violaxanthins, 4 vaucheriaxanthin esters, 2 phylloquinones, and 3 [4Fe–4S] clusters (Fig. S3, Table S2).

**Figure 1.**
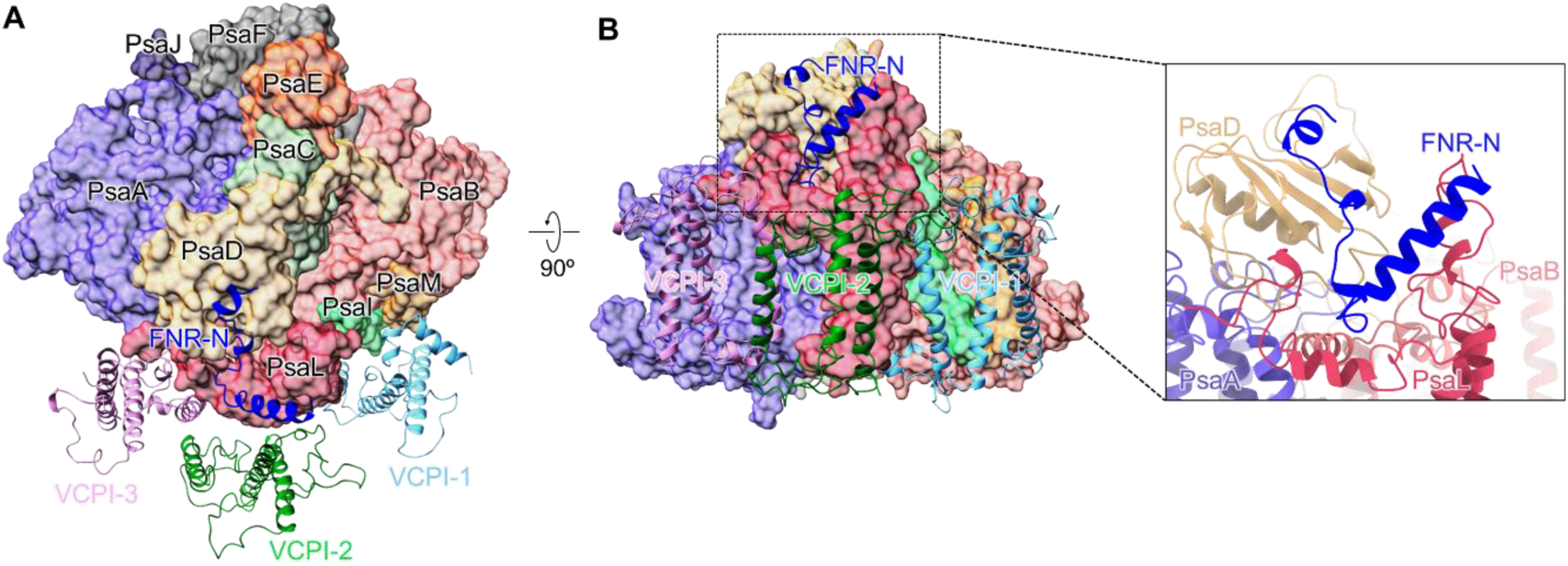
Overall structure of the eustigmatophyte PSI–VCPI supercomplex. **(A)** The PSI–VCPI supercomplex viewed from the stromal side. The core subunits are labeled by their name and the three VCPIs are labeled VCPI-1, VCPI-2 and VCPI-3. **(B)** Side view of the PSI–VCPI supercomplex. The boxed region is enlarged to show the binding site of FNR-N.

### Structure of the PSI core

The PSI core is composed of 10 resolved subunits: PsaA-F, PsaI, PsaJ, PsaL, and PsaM (Figs. 1, S4). Notably, the structures of PsaL and PsaD are compatible with the docking of the N-terminus of FNR (FNR-N) (Fig. 2, Fig. S5). VsPsaL contains an approximately 30-residue insertion (E105–D136) that extends across the stromal surface and contacts FNR-N through hydrogen-bonding and hydrophobic interactions (Figs. 2A, 2B). A corresponding PsaL insertion participates in FNR docking in *N. oceanica*, and sequence comparisons indicate that PsaL is conserved among sampled eustigmatophytes (Fig. S6) (31). The N-terminal region of VsPsaD adopts a lineage-specific conformation and provides additional contacts with FNR-N (Figs. 2C, D and S7).

**Figure 2.**
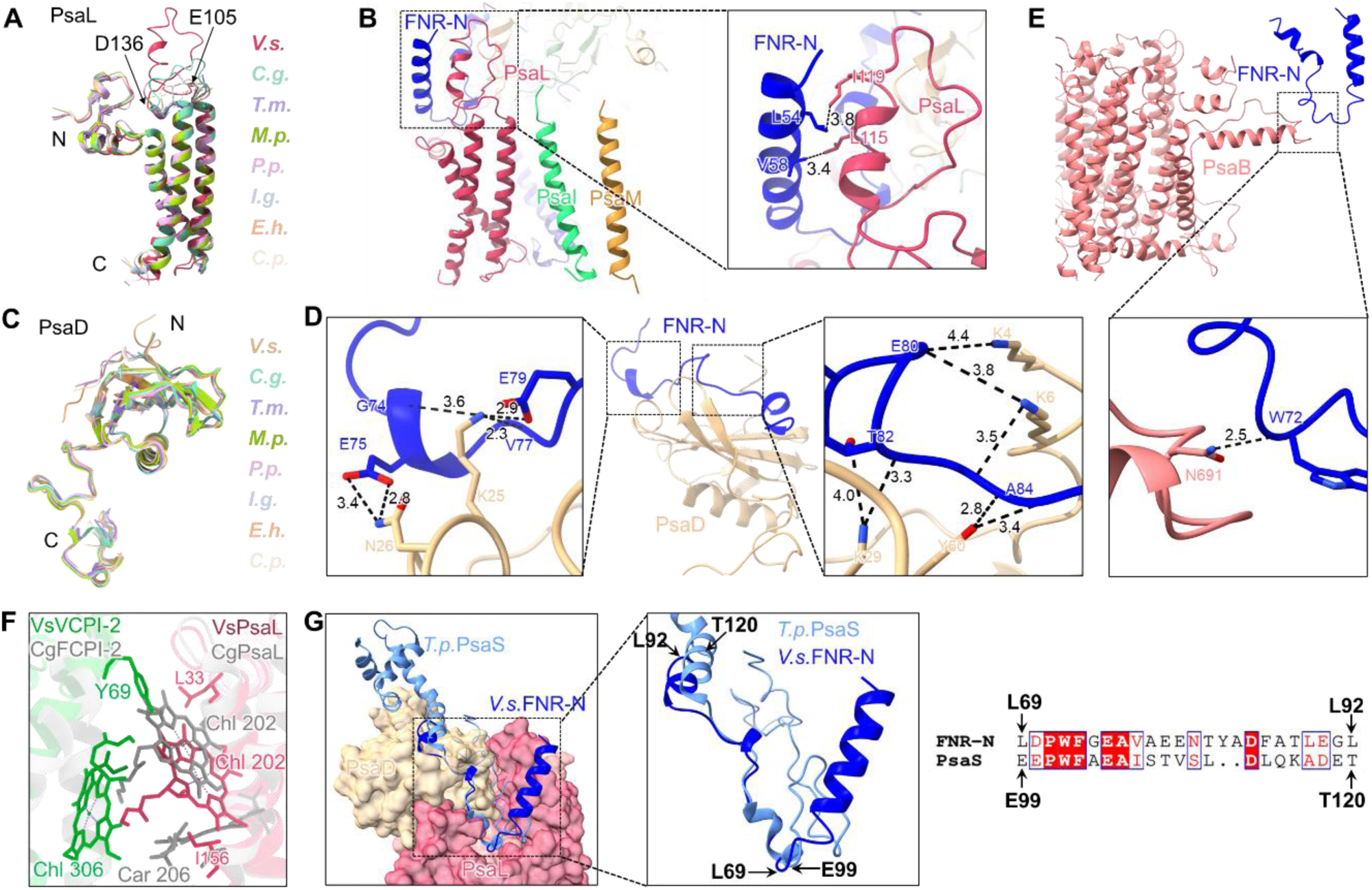
Structural basis for the association of FNR-N with the PSI core in the eustigmatophyte VsPSI– VCPI. **(A)** Structural comparison of PsaL from *Vischeria stellata* (*V.s.*) with those from red-lineage algae. Diatom *Chaetoceros gracilis* (*C.g.*), xanthophyte *Tribonema minus* (*T.m.*), phaeophyte *Macrocystis pyrifera* (*M.p.*), red alga *Porphyridium purpureum* (*P.p.*), haptophyte *Isochrysis galbana* (*I.g.*), coccolithophore *Emiliania huxleyi* (*E.h.*) and cryptophyte *Chroomonas placoidea* (*C.p.*). **(B)** Interactions between FNR-N and the extended loop of PsaL. Squared area is enlarged, showing the detailed hydrogen-bonding interactions. Interactions are indicated by black dashed lines with distances labeled in Å. **(C)** Structural comparison of PsaD from *V. stellata* (*V.s.*) with those from red-lineage algae. **(D)** Interactions between FNR-N and the N-terminus of PsaD. Squared areas are enlarged, showing the detailed hydrogen-bonding interactions. Interactions are indicated by black dashed lines with distances labeled in Å. **(E)** Interactions between FNR-N and the PsaB. Squared area is enlarged, showing the detailed hydrogen-bonding interactions. Interactions are indicated by black dashed lines with distances labeled in Å. **(F)** Comparison of the binding position and microenvironment of pigments in VsPsaL and diatom CgPsaL. **(G)** Comparison of the locations and structures of FNR-N from *V. stellata* (*V.s.*) and PsaS from *Thalassiosira pseudonana* (*T.p.*). Squared area is enlarged, showing the structural similarity between FNR-N L69-L92 and PsaS E99-T120.

Compared with the PSI cores of xanthophytes and diatoms within Ochrophyta (14, 18), the VsPSI core lacks PsaR and PsaS (Fig. S4). Although the *V. stellata* transcriptome contains a gene encoding PsaR, no density was detected at its expected binding site, suggesting that PsaR may have dissociated during purification or is substoichiometrically associated with the complex. In contrast, neither PsaS density nor a corresponding transcript was identified, suggesting the absence of PsaS from VsPSI–VCPI.

The PSI core binds 83 Chl *a*, 14 *β-*Car and 1 Vio molecules (Table S2). Several peripheral pigments near PsaB and PsaF were not assigned because of the limited map densities. In contrast to the PSI cores of other red-lineage algae, the VsPSI core lacks the conserved PsaL-Car 206 and exhibits a pronounced shift of the conserved PsaL-Chl 202 (Figs. 2F, S8). The shifting of PsaL-Chl 202 arises from altered microenvironment at the binding interface between VCPI-2 and the PSI core (Fig. 2F). Structural rearrangements and the altered docking position of VCPI-2 bring Y69 and Chl 306 into steric conflict with Chl 202. In addition, VsPsaL-L33 prevents Chl 202 from occupying its conserved canonical binding site, while the repositioned Chl 202, together with VsPsaL-I156, sterically precludes the binding of Car 206.

### An FNR N-terminal segment binds the stromal PSI surface

An additional V-shaped density was observed on the stromal surface of VsPSI–VCPI, adjacent to PsaB, PsaD, and PsaL (Figs. 1, 2). A 43-residue segment corresponding to residues 50–92 of the transcriptome-derived FNR sequence could be modeled into this density (hereafter designated FNR-N). FNR-N occupies a position analogous to the N-terminal extension of FNR in the recently reported *N. oceanica* PSI–VCP–FNR structure (31), indicating that this docking arrangement is conserved across Eustigmatophyceae. This FNR-N domain is absent from the previously reported FNR structures (33, 34), and sequence analysis indicates that this domain exists only in eustigmatophytes and is highly conserved (Fig. S5). In VsPSI–VCPI, FNR-N interacts primarily with the eustigmatophyte-conserved insertion in PsaL and the N-terminal region of PsaD. No interpretable density was detected for the catalytic FAD- and NADP(H)-binding regions of FNR, suggesting remarkable conformational flexibility of the intact enzyme relative to the PSI core; however, partial occupancy or dissociation during purification cannot be excluded. Consequently, the present structure defines the FNR-docking segment but neither resolves the position of the catalytic domain nor establishes how Fd engages PSI-associated FNR. Direct association of FNR with PSI may retain the mobile catalytic domain near the PSI acceptor side, potentially facilitating sequential electron transfer through soluble Fd while preserving the conventional PSI–Fd–FNR pathway.

In diatom and xanthophyte PSI–LHCI structures, part of the site occupied by FNR-N corresponds to the binding position of PsaS (Fig. 2G) (17, 18). A short segment of FNR-N (L69–L92) and a region of PsaS from *Thalassiosira pseudonana* (E99–T120) adopt similar secondary structures at the PSI stromal surface. This correspondence raises two possibilities: the proteins may share a remote evolutionary relationship, or they may have convergently evolved to interact with the same PSI region. Distinguishing between these possibilities will require broader phylogenetic sampling and structure-guided sequence analysis. The overlapping positions also raise the hypothesis that PsaS influences the organization of the PSI acceptor-side surface, although its direct involvement in Fd or FNR binding remains unproven.

### Conservation and divergence of FNR docking in eustigmatophytes

The VsPSI–VCPI structure provides an independent terrestrial counterpart to the marine *N. oceanica* PSI–VCP– FNR complex (31). Both structures resolve an FNR N-terminal extension at the PsaB–PsaD–PsaL region and identify an approximately 30-residue stromal insertion in PsaL as a principal docking element. However, the structures differ in both interface composition and structural order. The NoPSI–VCP–FNR interface includes an additional contact with a specialized C-terminal extension of PsaI, whereas no corresponding PsaI–FNR contact is resolved in *V. stellata*. Furthermore, NoPSI–VCP–FNR retains weak density for the mobile catalytic FNR domain, whereas VsPSI–VCPI resolves only the 43-residue docking segment. The most striking difference is antenna organization: VsPSI–VCPI contains three VCPIs confined to the PsaL–PsaI–PsaM side, whereas nine VCPs are distributed more extensively around NoPSI–VCP–FNR. Thus, the FNR-tethering mechanism is retained despite substantial diversification in the size and architecture of the eustigmatophyte PSI antenna.

### Structure and organization of VCPIs

Each VCPI subunit contains three transmembrane helices, consistent with the conserved architectures of previously reported LHCs from algae and land plants (Figs. 3A-3C, S9). Phylogenetic analysis assigned the three VCPI subunits to different antenna families: VCPI-1 to RedCAP, VCPI-2 to Lhcf, VCPI-3 to a CgLhcr9-like group (Fig. S10).

**Figure 3.**
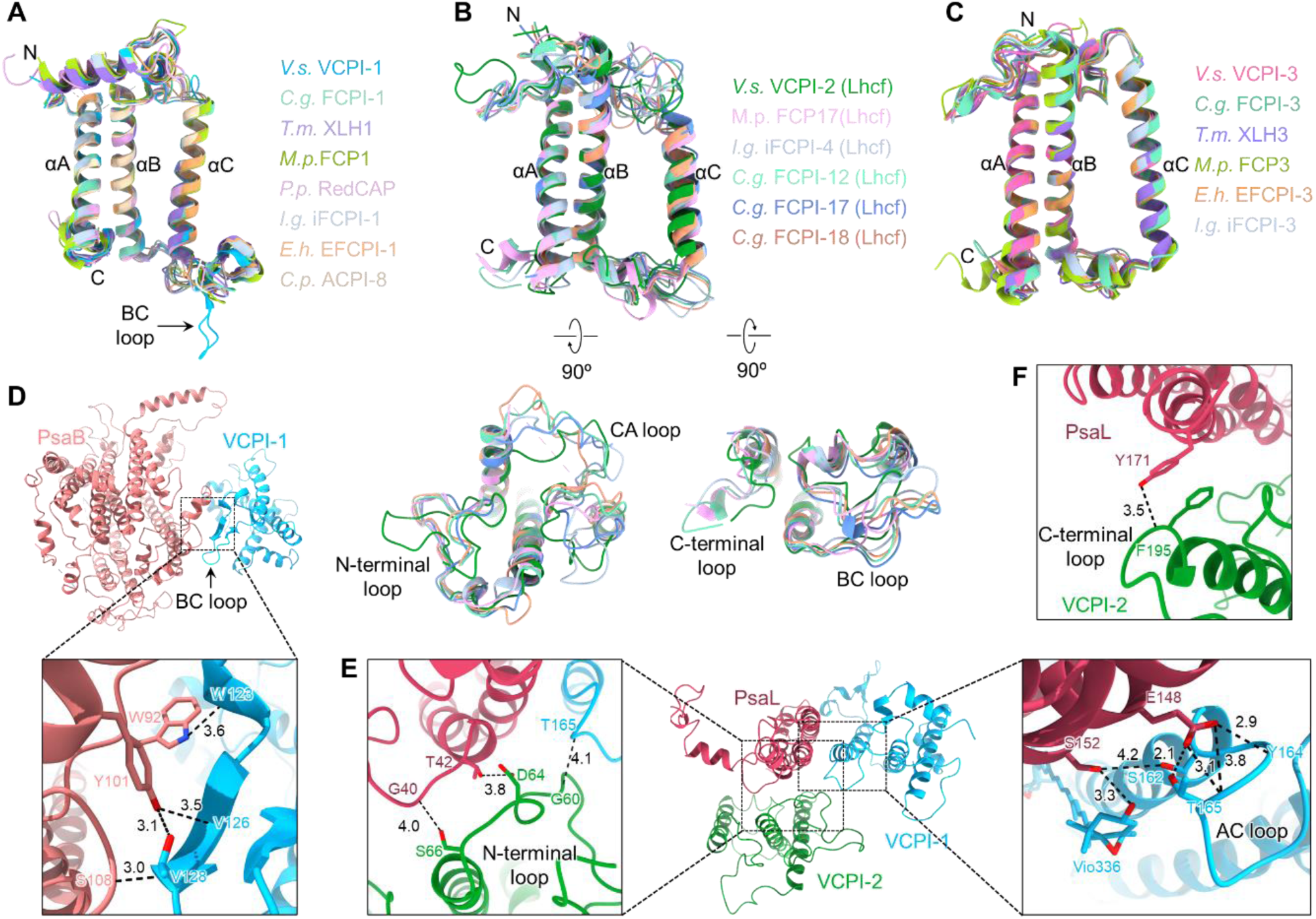
Structural characteristics and inter-subunit interactions of VCPIs in eustigmatophyte VsPSI– VCPI. **(A)** Structural comparison of RedCAPs from *V. stellata* (*V.s.*), diatom *Chaetoceros gracilis* (*C.g.*), xanthophyte *Tribonema minus* (*T.m.*), phaeophyte *Macrocystis pyrifera* (*M.p.*), red alga *Porphyridium purpureum* (*P.p.*), haptophyte *Isochrysis galbana* (*I.g.*), coccolithophore *Emiliania huxleyi* (*E.h.*) and cryptophyte *Chroomonas placoidea* (*C.p.*). The N-terminus, C-terminus, and three transmembrane helices are labeled as N, C, αA, αB, and αC, respectively. **(B)** Structural comparison of VCPI-2 from *V. stellata* (*V.s.*) with Lhcf-type antennas from PSI–LHCI of *Macrocystis pyrifera* (*M.p.*), *Isochrysis galbana* (*I.g.*) and *Chaetoceros gracilis* (*C.g.*). **(C)** Structural comparison of VCPI-3 from *V. stellata* (*V.s)* with its counterparts from red-lineage algae. **(D)** Interactions between the BC loop of VCPI-1 and PsaB. Squared area is enlarged, showing the detailed hydrogen-bonding interactions. Interactions are indicated by black dashed lines with distances labeled in Å. **(E)** Stromal-side interactions among AC loop of VCPI-1, N-terminal loop of VCPI-2, and PsaL. Squared area is enlarged, showing the detailed hydrogen-bonding interactions. Interactions are indicated by black dashed lines with distances labeled in Å. **(F)** Lumenal-side Interactions between C-terminal loop of VCPI-2 and PsaL. Interactions are indicated by black dashed lines with distances labeled in Å.

Homologs belonging to the same clades as VCPI-1 and VCPI-3 exist in PSI–LHCI from diatoms, xanthophytes, phaeophytes, and haptophytes (Fig. S10). They exhibit similar binding positions, structures, and sequences, indicating their close evolutionary relationship (Figs. S11, S12). Although sharing a similar overall architecture with RedCAP in other algae, VCPI-1 possesses several specialized loop structures. Its BC loop extends further toward the lumenal side and forms additional interactions with PsaB (Figs. 3A, 3D, S12A), whereas its AC loop adopts a different conformation, forming interactions with PsaL and VCPI-2 on the stromal side (Fig. 3E). By contrast, compared with diatom FCPI-2, xanthophyte XLH2, phaeophyte FCP2, and haptophyte iFCPI-2/EFCPI-2, VCPI-2 shifts toward VCPI-3 and undergoes rotational rearrangement (Fig. S11). In addition, VCPI-2 is absent in marine *N. oceanica* PSI–VCP–FNR (31). Phylogenetic analysis showed that VCPI-2 does not cluster with Lhcq-type FCPI-2/XLH2 and Lhcr-type iFCPI-2, but instead groups with Lhcf-type antenna proteins (Fig. S10). Together with the structural variability observed at this position (Fig. S11), this finding indicates that the identity and orientation of the corresponding antenna subunit are not conserved across these lineages and may contribute to lineage-specific PSI–LHCI assembly. Although the N-terminal and C-terminal loops are relatively conserved among these Lhcf-type subunits, their conformations differ markedly in VCPI-2, corresponding to its distinct binding position in VsPSI–VCPI (Fig. 3B). Its N-terminal loop contacts PsaL and VCPI-1 on the stromal side, while its C-terminal loop interacts with PsaL on the lumenal side (Figs. 3E and 3F).

Unlike the multilayered or ring-like LHCI arrangements in other red-lineage algae, the three VCPIs in VsPSI– VCPI form a compact belt along the PsaL–PsaI–PsaM side of the PSI core, representing the smallest antenna reported among structurally characterized red-lineage PSI–LHCI supercomplexes. The reduced antenna may lead to a smaller PSI light-harvesting cross-section and could be relevant to adaptation to high irradiance in terrestrial habitats. However, comparative physiological measurements and analysis of native complexes under different irradiances are needed to determine whether the three-VCPI organization represents a constitutive terrestrial feature, a condition-dependent assembly state, or partial antenna loss during purification.

### Pigment arrangement in VCPIs

The three VCPI subunits contain 31 Chl a, 8 Vio, and 4 Vau -ester molecules (Table S2). Most Chl-binding sites in VCPIs are conserved with previously reported red-lineage LHCIs, whereas several specific pigment-binding sites were identified in VCPIs (Table S3) (5, 8, 11, 13, 14, 18, 19).

Nine Chl-binding sites were identified in RedCAP-type VCPI-1 (Fig. 4A). Among them, Chls 303 to 307, 310, 311, and 313 are conserved in previously reported RedCAP antennae (Fig. S13A). By contrast, Chl 312 is absent from all previously reported RedCAP antennae as well as from the RedCAP antenna of marine *N. oceanica* (31) and binds to the lumenal side of VCPI-1 αA helix (Fig. 4A). It locates at the interface between VCPI-1 and PsaM and lies in close proximity to PsaM/Car101, which may facilitate energy transfer from antennae to the PSI core.

**Figure 4.**
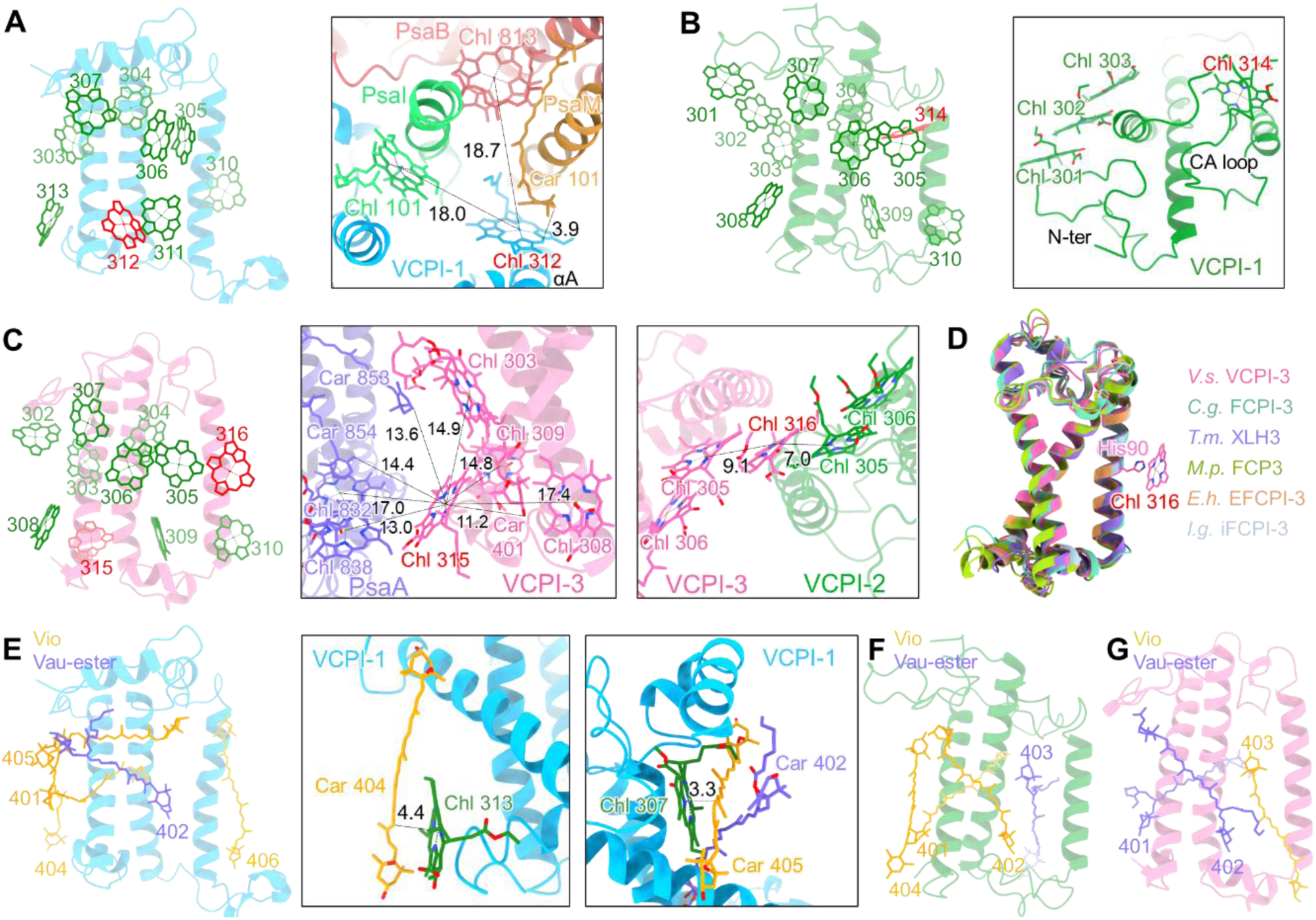
Pigment-binding sites in *V. stellata* VCPIs. **(A)** Locations of the 9 Chls in VCPI-1 (left panel, conserved Chl sites in red-lineage algae are colored green) and the distance between the specific Chl 312 and neighboring pigments (right panel). Distances are indicated by black dashed lines and labeled in Å. **(B)** Locations of the 11 Chls in VCPI-2 (left panel, conserved Chl sites in red-lineage algae are colored green). The parallel arrangement of Chls 301, 302 and 303 and the location of the specific Chl 314 (right panel). **(C)** Locations of the 11 Chls in VCPI-3 (left panel, conserved Chl sites in red-lineage algae are colored green) and the distance between the specific Chl 315 and neighboring pigments (middle panel), and between the specific Chl 316 and neighboring pigments (right panel). Distances are indicated by black dashed lines and labeled in Å. **(D)** Structural comparison of VCPI-3 from *V. stellata* (*V.s.)* with its counterparts from *Chaetoceros gracilis* (*C.g.*), *Tribonema minus* (*T.m.*), *Macrocystis pyrifera* (*M.p.*), *Emiliania huxleyi* (*E.h.*), and *Isochrysis galbana* (*I.g.*). Chl 316 and its coordinating residue His90 in VCPI-3 are shown. **(E)** Locations of the 5 Cars in VCPI-1 (left panel) and the positions of Car 404 (middle panel) and Car 405 (right panel) relative to neighboring pigments. Distances are indicated by black dashed lines and labeled in Å. **(F)** Locations of the 4 Cars in VCPI-2. **(G)** Locations of the 3 Cars in VCPI-3.

Eleven Chl-binding sites were identified in Lhcf-type VCPI-2 (Fig. 4B). Among them, Chls 302 to 310 are highly conserved in red-lineage algal antennae (Fig. S13B). Chl 301 is located near the N-terminal loop of VCPI-2, and no corresponding site has been found in reported Lhcf-type antennae (13, 14, 19) (Fig. 4B). Chl 301 is arranged parallel to Chls 302 and 303, potentially facilitating EET among them. In addition, the specific Chl 314 is located near the αC helix and is associated with the CA loop of VCPI-2 (Fig. 4B).

CgLhcr9-like VCPI-3 coordinates 11 Chls. Chls 302 to 310 are conserved sites, whereas Chls 315 and 316 are specific sites to VCPI-3 and is absent from all previously reported CgLhcr9-like antennae including that of marine *N. oceanica* (Figs. 4C, S13C) (31). Chl 315 binds to the C-terminal loop and forms close interactions with pigments in PsaA and VCPI-3, potentially facilitating EET from antennae to the PSI core (Fig. 4C). Chl 316 is located near the αC helix and coordinated by His90. His90 of VCPI-3 is substituted by Gly in the corresponding antennae of diatoms, xanthophytes, phaeophytes, and haptophytes, leading to the absence of Chl 316 (Fig. 4D). Chl 316 is positioned at the interface between VCPI-3 and VCPI-2, and is arranged parallel to red Chl 305/306 pairs of VCPI-2 and VCPI-3, suggesting its potential role in facilitating EET between VCPI-2 and VCPI-3 (Fig. 4C).

All three VCPI subunits bind two types of Cars: Vio and Vau -ester. VCPI-1 contains four Vio and one Vau-ester molecules, among which Vio401, Vau-ester402, and Vio406 are conserved Car-binding sites in RedCAP antennae (Figs. 4E, S13B). Vio404 resides near the αC helix and C-terminal loop, while Vio405 is parallel to the N-terminal helix and is sandwiched between VCPI-1/Chl 307 and VCPI-1/Vau-ester402 (Fig. 4E). Except for the detection of the Car 405 site in xanthophyte RedCAP antenna and Car 404/405 sites in RedCAP antenna of marine *N. oceanica*, neither Car 405 nor Car 404 have been identified in previously reported RedCAP antennae (Fig. S13B). They are parallel to the chlorin ring of Chl and in close proximity to Chl at a distance of ∼ 4 Å, enabling the overlap of their π-electron clouds with those of Chls (Fig. 4E). This may facilitate EET from Chl to Vio, suggesting their potential roles in dissipating excess excitation energy under high light of soil habitat. VCPI-2 contains three Vio and one Vau-ester molecules, whereas VCPI-3 contains one Vio and two Vau-ester molecules, and all these pigment-binding sites are conserved (Figs. 4F-4G, S13D, S13F).

In summary, each VCPI contains conserved and lineage-specific pigment sites that may form excitation-transfer routes within the compact antenna, although their roles in energy transfer and dissipation require spectroscopic and light-dependent validation. The lineage-specific pigment sites Chls 312/315/316 and pigments in VCPI-2 which are absent in marine *N. oceanica* due to the missing of corresponding antenna constitute a pigment organization that is markedly different from that of marine *N. oceanica*, which may reflect the adaptation strategy of energy transfer in soil VsPSI–VCPI under terrestrial environment.

### Energy transfer within VsPSI–VCPI

The distinctive position of VCPI-2 and the additional pigment-binding sites identified in the three antennae generate a pigment network that differs from those of previously reported PSI–LHCI complexes (8, 11, 13, 14, 18, 19). We calculated pairwise EET rates between Chls within a Förster framework and estimated energy transfer between pigment groups using generalized Förster (GF) theory (Fig. 5) (35, 36). To minimize structural-model uncertainty, before performing the energy-transfer calculations, we optimized the structure of the entire complex through molecular dynamics simulations, refined the pigment geometries using a QM/MM approach, and corrected some portions of the protein periphery in contact with the membrane with reference to the crystallographic structure. The calculations indicate strongly coupled neighboring Chls and candidate transfer routes from each VCPI to the PSI core. Sub-picosecond transfer times indicate strong pigment coupling but should not be interpreted quantitatively because the incoherent Förster approximation becomes unreliable in this regime.

**Figure 5.**
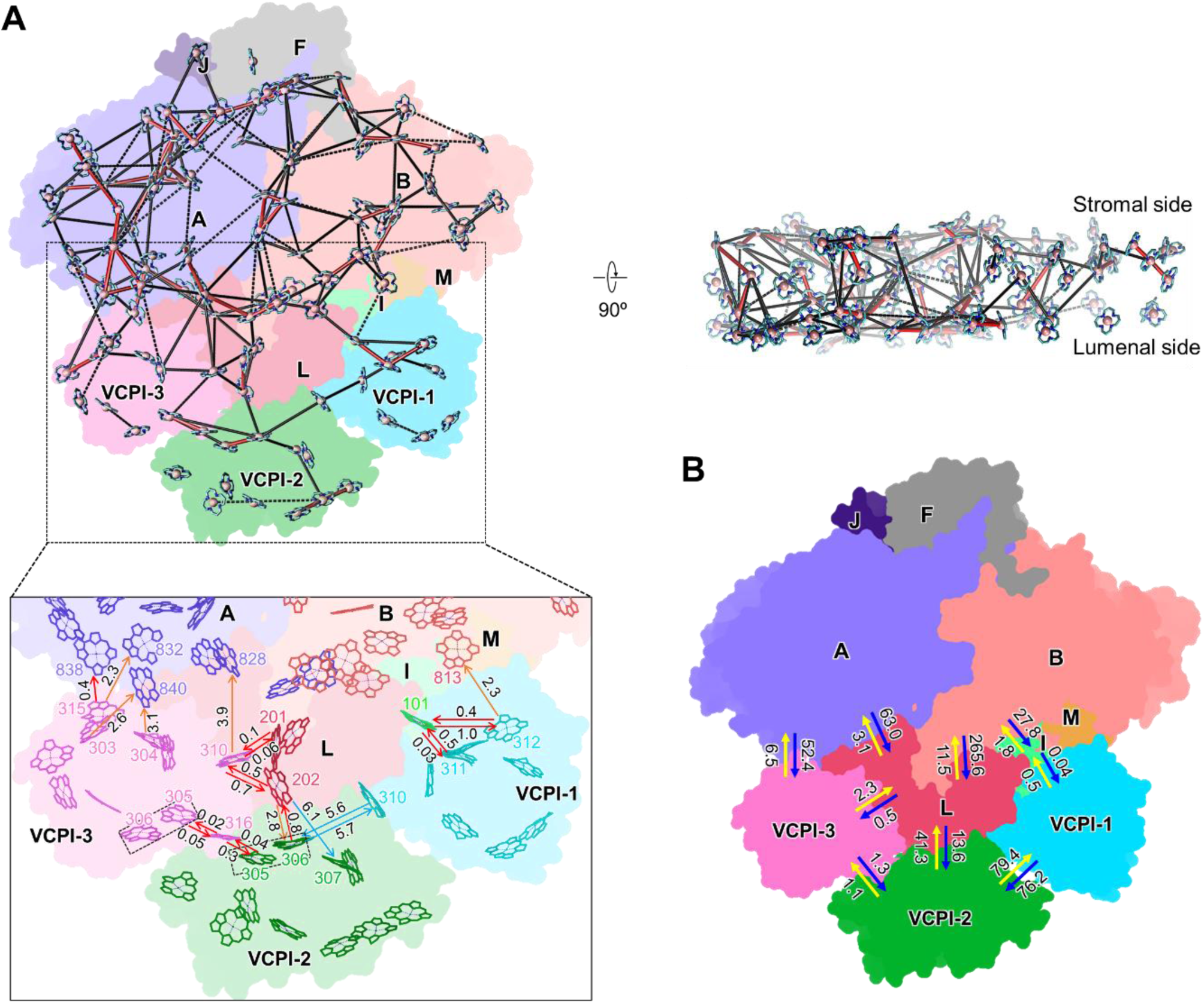
Excitation energy transfer in eustigmatophyte VsPSI–VCPI. **(A)** The inter-pigment energy transfer network generated by quantum simulation based on Förster. In the upper panels, the lines indicate rates faster than 1 ps (red), in the 1-10 ps range (black solid lines) and in the 10-20 ps range (black dashed lines). Inter-pigment rates slower than 20 ps are omitted. The enlarged view at the lower panel indicates the key energy transfer pathways and Chls. Energy transfer rates faster than 2 ps (red arrow), between 2 and 5 ps (orange arrow), and between 5 and 10 ps (blue arrow). The red Chl 305/306 pairs are indicated by dashed box. **(B)** Energy transfer rates labeled in ps between VCPIs and between VCPIs and PSI core subunits simulated based on generalized Förster theory.

GF simulations reveal that VCPI-1 can efficiently transfer energy to PsaB via the mediation of PsaI (2.3 ps) (Fig. 5B). Chl 311 and the specific Chl 312 of VCPI-1 perform pivotal functions, enabling ultrafast energy transfer to PsaI-Chl 101 (∼0.5 ps) (Fig. 5A). Chl 312 can also directly deliver energy efficiently to PsaB-Chl 813 (2.3 ps), which accelerates EET from VCPI-1 toward PSI. VCPI-2 routes excitation energy to PSI mainly through VCPI-3: VCPI-2–VCPI-3–PsaA (7.6 ps), VCPI-2–VCPI-3–PsaL–PsaA (6.5 ps), and VCPI-2–VCPI-3– PsaL–PsaB (14.9 ps) (Fig. 5B). The specific Chl 316 of VCPI-3 and Chl 305/306 pairs play critical roles in EET.

Chl 305/306 pair of VCPI-2 efficiently transfer EET to Chl 316 of VCPI-3 (0.04 ps) and subsequently to Chl 305/306 pair of VCPI-3 (0.02 ps) (Fig. 5A). VCPI-3 can transfer energy directly to PsaA (6.5 ps). It can also mediate EET to PsaA (5.4 ps) and PsaB (13.8 ps) via PsaL (Fig. 5B). Its Chls 303/304/310/315 play key roles in these energy transfer pathways (Fig. 5A). Chls 303/304 form efficient EET routes coupled with PsaA-Chl 840 (∼3.0 ps). Chl 310 transfers energy rapidly to PsaA-Chls 828 (3.9 ps) and PsaL-Chls 201/202 (0.06 ps/0.7 ps). The specific Chl 315 further reinforces EET by establishing additional high-efficiency pathways with PsaA-Chls 832/838 (2.3 ps/0.4 ps).

Conserved pigments, such as the Chl 305/306 pair and Chls 303/304/310/311, dominate energy transfer processes, similar to those reported in red-lineage PSI–LHCIs (8, 11, 13, 20). Moreover, the species-specific Chls 312/315/316 within eustigmatophyte VsVCPI also form efficient EET pathways with surrounding pigments, make substantial contributions to the energy transfer within VsPSI–VCPI which are absent in marine *N. oceanica* PSI–VCPI. This specialized pigment configuration may act as a compensatory mechanism to offset the limited light-harvesting capacity of its simplified antenna system.

The calculations also predict slower net transfer from the core to peripheral modules, including PsaA to VCPI-3 (52.4 ps), PsaA to PsaL (63.0 ps), PsaB to PsaL (265.6 ps), and PsaB to PsaI (27.8 ps) (Fig. 5B). These transfer rates are 10–20-fold slower than those of forward energy transfer from peripheral subunits to the PSI core, confirming that energy flow toward the PSI core holds an absolute advantage, consistent with energetically favored core-directed EET. Intriguingly, the PsaI and PsaL subunits, which mediate EET from light-harvesting antennae to the PSI core, can also transfer energy efficiently back to the antennae: PsaI to VCPI-1 (0.04 ps), PsaL to VCPI-2 (13.6 ps), and PsaL to VCPI-3 (0.5 ps) (Fig. 5B). This indicates that PsaI and PsaL can rapidly shunt excess excitation energy back to light-harvesting antennae for thermal dissipation, restricting excitation energy passing into the PSI core to protect it from photodamage. PsaI efficiently transfers energy to Chl 311 (0.03 ps) and the specific Chl 312 (1.0 ps) of VCPI-1, whereas PsaL rapidly delivers energy to Chls 306/307 of VCPI-2 (2.8 ps/6.1 ps) and Chl 310 of VCPI-3 (0.1 ps/0.5 ps) (Fig. 5A). These pairwise rates indicate bidirectional energetic connectivity but do not alone establish net back-transfer or thermal dissipation. Two specific Cars (Vio 404 and Vio 405) tightly associated with Chls in VCPI-1 are proposed to contribute to energy dissipation. These energy-transfer features likely represent a photoprotective strategy evolved by VsPSI–VCPI to adapt to high irradiance in soil habitats. The involvement of specific pigments Chl 312, Vio 404, and Vio 405 imply that VCPI-1 may play a critical role in energy dissipation.

### Insights into the evolution and adaptation of eustigmatophyte VsPSI–VCPI

Within the phylogenetic framework based on nuclear genomes, the Ochrophyta is generally divided into two major clades: Diatomista and Chrysista (26, 37). Diatoms fall into the former group, whereas Eustigmatophyceae and RPX clade comprising Raphidophyceae, Phaeophyceae, and Xanthophyceae belong to the latter. This indicates that eustigmatophytes and diatoms are not closely-related sister groups; instead, they underwent independent evolution along distinct trajectories following the early divergence of the Ochrophyta. Eustigmatophyceae is widely regarded as an independent evolutionary lineage which is more closely related to the RPX clade and shares a common ancestor with RPX clade (26).

Cryptophytes originated from secondary endosymbiosis with a red alga (38). Their PSI associate with two types of light-harvesting antennae, Lhcr and RedCAP, consistent with those of red algae, and their light-harvesting antenna systems share a similar organization with those of red algae (Fig. 6). This indicates that their PSI–LHCI complexes derive from red algal PSI–LHCI. Nuclear genomic analyses suggest that ochrophytes arose via tertiary endosymbiosis with a cryptophyte (39). In addition to Lhcr and RedCAP, ochrophyte LHCIs have further diversified into Lhcq and Lhcf isoforms (Fig. 6). Phylogenetic analyses demonstrate that the majority of their Lhcr-type LHCIs and RedCAP proteins have a close genetic relationship with those of cryptophytes, whereas a minor subset of Lhcr-type LHCIs trace their origin to red algae (40). The structures and arrangement of Lhcr-type LHCIs and RedCAP in ochrophytes resemble those found in cryptophytes and red algae, supporting their PSI–LHCI evolved from ancestral counterparts in cryptophytes and red algae.

**Figure 6.**
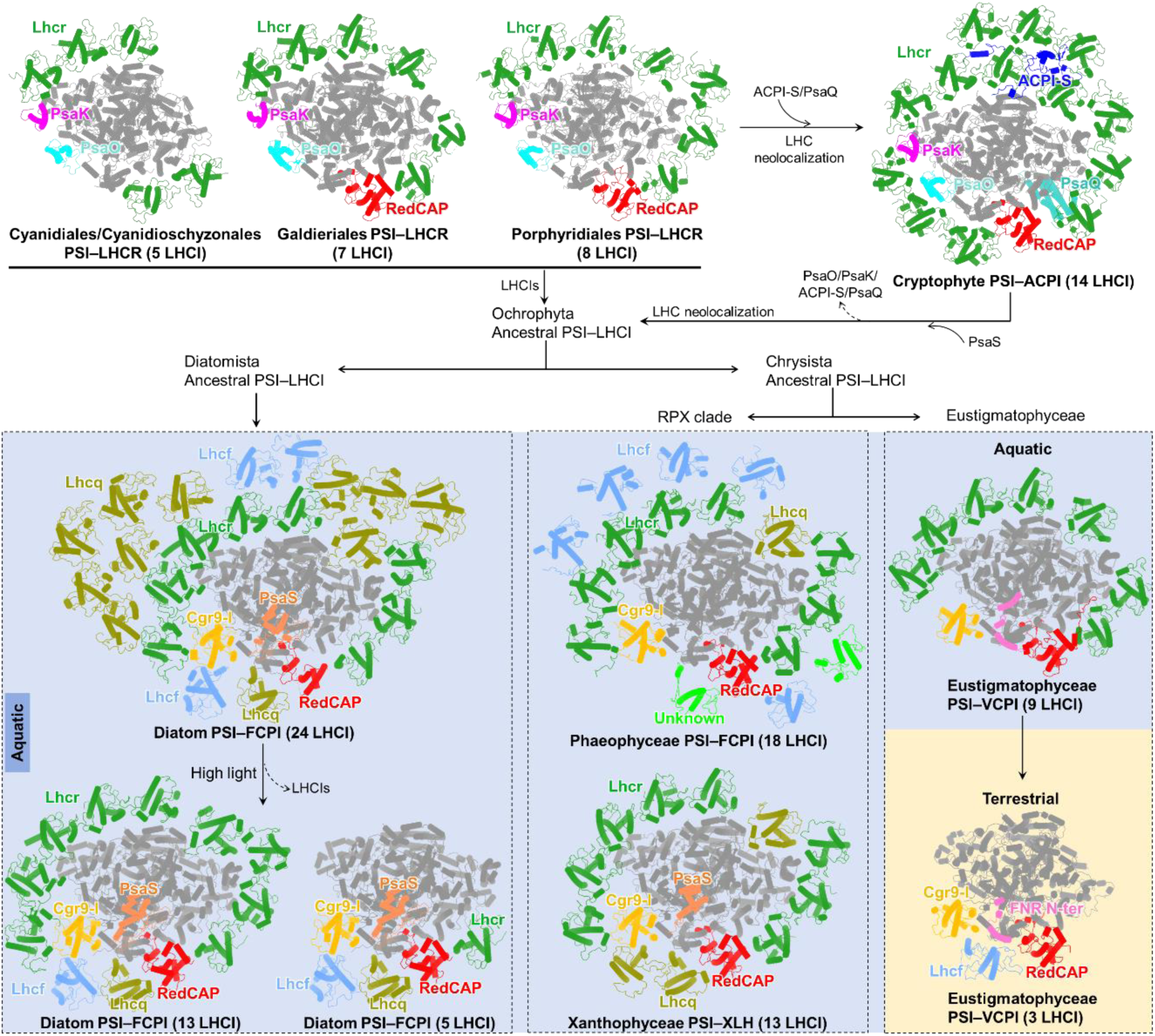
Insights into the evolution and adaptation of Ochrophyta PSI–LHCI supercomplexes. PDB IDs of PSI–LHCI structures: PSI–LHCR of red alga *Cyanidioschyzon merolae* (5ZGB); PSI–LHCR of red alga *Galdieria sulphuraria* (9KC5); PSI–LHCR of red alga *Porphyridium purpureum* (7Y5E); PSI–ACPI of cryptophyte *Chroomonas placoidea* (7Y7B); PSI–24FCPI of diatom *Chaetoceros gracilis* (6LY5); PSI–13FCPI of diatom *Thalassiosira pseudonana* (8ZEH); PSI–5FCPI of diatom *Thalassiosira pseudonana* (8ZET); PSI– FCPI of phaeophyte *Macrocystis pyrifera* (9YGV); PSI–XLH of xanthophyte *Tribonema minus* (9M4F); PSI– VCPI of eustigmatophyte *Nannochloropsis oceanica* (8ZO9). Cgr9-l is abbreviation of CgLhcr9-like.

Ochrophyte lineages exhibit substantial variation in PSI antenna organization (Fig. 6). Diatoms are dominant marine photosynthetic organisms featuring a drastically expanded PSI antenna system with as many as 24 light-harvesting subunits (14), an expanded arrangement consistent with efficient light harvesting in fluctuating marine environments. Giant kelp (Phaeophyceae) constitutes critical components of coastal ecosystems, and kelp forests exhibit net primary productivity three times higher than marine phytoplankton, and its PSI associates with 18 antenna subunits (19). The xanthophyte *T. minus* inhabits relatively static shallow-water habitats and possesses a PSI bound to only 13 light-harvesting antennae (18), fewer than diatoms and giant kelp. This structural feature likely underpins its adaptation to high-light shallow-water environments. Similarly, diatom PSI also contains a reduced number of light-harvesting antennas (13 or 5 subunits) under high-light conditions (17).

VsPSI–VCPI contains only three peripheral antennae, substantially fewer than the nine VCPs reported in marine *N. oceanica* and the larger antenna systems of other structurally characterized ochrophyte PSI–LHCI (14, 18, 19). The minimal VsPSI–VCPI antenna system may constitute an evolutionary adaptation to high-light environment on the soil surface, restricting the influx of excess light energy into PSI. Analogously, during the evolutionary transition from aquatic green algal PSI–LHCI to land plants PSI–LHCI, the number of light-harvesting antennae declined from ten to four as an adaptation to the high-irradiance environment of terrestrial habitats (41–44). Furthermore, previous studies reported that the green alga *Chlorella ohadii*, which dwells in desert soil biological crusts, reduces its LHCI antenna system under high light to decrease the PSI light-harvesting cross-section and mitigate photodamage risks (24). Collectively, the reduction in the antenna system may represent a universal strategy adopted by photosynthetic organisms to cope with intense terrestrial light during aquatic-to-terrestrial evolutionary transitions. Whether the three-VCPI organization of *V. stellata* is constitutive or environmentally regulated to cope with intense terrestrial light during aquatic-to-terrestrial evolutionary transitions will require comparison of the structures and photosynthetic performance of VsPSI–VCPI complexes across light conditions.

Phylogenetic and structural analyses reveal that VCPI-1 and VCPI-3 share high sequence and structural homology with LHCIs occupying equivalent binding sites within the PSI–LHCIs of diatoms, xanthophytes and phaeophytes (Figs. S10, S11), indicating a shared evolutionary ancestor and supporting the evolutionary origin of eustigmatophytes. By contrast, VCPI-2 is classified as an Lhcf-type antenna, distinct from the LHCI at corresponding position in diatom, xanthophyte and phaeophyte PSI–LHCI (Fig. S10). Its structure and binding position have diverged substantially, generating antenna characteristics specific to eustigmatophytes, and matching the assembly of the simplified VsPSI–VCPI antenna system (Fig. S11). These observations suggest that the specialized organization of the VsPSI–VCPI antenna may contribute to light-energy conversion in its terrestrial soil habitat.

The occurrence of closely related FNR-N docking arrangements in *V. stellata* and *N. oceanica* indicates that PSI recruitment of FNR is not restricted to Nannochloropsis but is likely conserved more broadly within Eustigmatophyceae (Fig. 2). Both complexes use an extended FNR N-terminus and an inserted stromal region of PsaL, whereas the NoPSI–VCPI interface additionally involves a specialized PsaI extension. Thus, eustigmatophyte PSI appears to retain a common FNR-tethering principle, which may differ from the conventional Fd-dependent pathway. The tight interaction between FNR and the PSI core may accelerate electron transport, rapidly delivering abundant electrons generated under high light to NADP⁺, which may represent an adaptive mechanism of *V. stellata* to stressful soil habitats. On the other hand, because the catalytic FNR domain is unresolved in VsPSI–VCPI and electron-transfer activity was not measured, the present data do not indicate a pathway that bypasses Fd or establish enhanced linear electron flow. FNR-N may tether a mobile catalytic domain near the PSI acceptor side while allowing sequential interaction with soluble Fd.

Intriguingly, the N-terminus of eustigmatophyte FNR contains a region with high structural and sequence similarity to a region of PsaS from diatoms and xanthophytes (Fig. 2F), and both regions occupy identical binding sites on the stromal surface of PSI. This resemblance suggests that FNR-N may represent alternative lineage-specific solutions for organizing the PSI acceptor-side surface in eustigmatophyte. In contrast, PsaS may be involved in the binding of Fd and FNR proteins in diatoms and xanthophytes. However, positional and structural similarity alone cannot distinguish remote homology from convergent binding or establish a direct role for PsaS in recruiting Fd or FNR. Broader taxonomic sampling and structure-guided phylogenetic analysis will be required to test these hypotheses.

### Conclusions

The 2.44 Å cryo-EM structure of VsPSI–VCPI reveals an exceptionally compact antenna containing only three VCPI subunits and an FNR N-terminal segment tethered to the stromal surface of PSI. Comparison with *N. oceanica* PSI–VCPI indicates that FNR docking through lineage-specific elements of FNR, PsaL, and PsaD subunits is conserved despite extensive divergence in antenna size, composition, and organization. Structure-based computational calculations further indicate potential routes for excitation energy transfer and energetic connectivity between the compact antenna and the PSI core. Although the physiological consequences of antenna reduction and FNR tethering remain to be established, our findings suggest that PSI has evolved as a modular complex whose architecture can be extensively remodelled through changes in antenna composition without corresponding alterations to acceptor-side organization. This principle broadens our understanding of how photosynthetic machinery evolves across phylogenetic lineages and ecological transitions and generates testable hypotheses concerning the functional significance of antenna reduction and FNR tethering in terrestrial environments.

## Materials and Methods

### Purification of PSI–VCPI from *V. stellata*

The *Vischeria stellata* strain used in this study was obtained from the College of Life Science and Technology, Guangxi University, China. Cultures were maintained in BG-11 medium at 20 °C under continuous illumination of 40 μmol photons m⁻² s⁻¹ and supplied with constant aeration. After 4 d of growth, logarithmic-phase cells were collected by centrifugation at 6000 × g for 12 min. The cell pellets were washed twice with MES1 buffer (25 mM MES-NaOH, pH 6.5), resuspended in MES1 buffer supplemented with protease inhibitor, and disrupted using a low-temperature ultra-high-pressure continuous-flow cell disrupter at 1900 bar. Residual intact cells and coarse debris were separated from the homogenate by centrifugation at 3000 × g for 2 min, after which the membrane fraction was recovered from the supernatant by centrifugation at 21,000 × g for 20 min. The membrane pellet was washed once with MES3 buffer (25 mM MES-NaOH, pH 6.5, 1.0 M betaine, and 1 mM EDTA), recovered again at 21,000 × g for 20 min, and finally suspended in MES4 buffer (25 mM MES-NaOH, pH 6.5, 1.0 M betaine, 10 mM NaCl, and 5.0 mM CaCl_2_). Membrane solubilization was carried out in MES4 buffer with 2.5% (w/v) n-dodecyl-α-D-maltopyranoside (α-DDM) at a final Chl a concentration of 0.5 mg mL⁻¹. The suspension was incubated on ice for 15 min and gently mixed at 5-min intervals. Insoluble material was removed by centrifugation at 21,000 × g for 15 min. The clarified extract was fractionated on a 10–30% discontinuous sucrose density gradient containing 0.02% α-DDM, with successive layers differing by 2% sucrose. Ultracentrifugation was performed at 230,500 × g for 20 h in an SW40 rotor. The green band corresponding to PSI–VCPI was collected and concentrated with a 100-kDa molecular weight cutoff filter (Amicon Ultra; Millipore). Throughout purification, samples were handled at 4 °C and dim light.

### Biochemical characterization of *V. stellata* PSI–VCPI

The absorption spectrum of purified PSI–VCPI was measured at room temperature using a Shimadzu UV–Vis 1900 spectrophotometer over the wavelength range of 400–750 nm. The subunit composition of PSI–VCPI was examined by SDS-PAGE using 8–16% gradient gels. Protein bands were excised from the gel, reduced, alkylated, and digested with trypsin. The resulting peptides were analyzed by LC–MS/MS using a nanoElute nano-flow LC system coupled to a timsTOF Pro mass spectrometer (Bruker). Peptides were separated on a Thermo Scientific EASY C18 analytical column (25 cm length, 75 μm inner diameter, 1.9 μm particle size). The raw MS data were searched against the selected protein database using MaxQuant 1.6.14.

The pigment composition of PSI–VCPI was assessed by HPLC following previously described procedures (13). Pigments were extracted from the purified complex with precooled 100% acetone overnight at 4 °C in dark and then applied to a C18 reversed-phase column (Waters). The eluates were monitored at 440 nm, and absorption spectra were recorded over a wavelength range of 300–800 nm. Four pigments, including Chl *a*, violaxanthin, vaucheriaxanthin ester, and β-carotene, were assigned on the basis of their characteristic absorption features and elution profiles, consistent with previous pigment profiles reported for *V. stellata* (45).

### Sequence analysis of *V. stellata* PSI–VCPI

*V. stellata* transcriptome was generated by Huada using the DNBSEQ platform. Total RNA was extracted from *V. stellata* cells and used for cDNA library construction. After quality filtering, the clean reads were assembled de novo, and coding sequences were predicted from the assembled transcriptome. Sequences of the PSI core subunits and VCPIs were identified by searching the transcriptome assembly with homologous sequences as queries. Sequence comparisons were performed using CLC Sequence Viewer 8.0 and displayed with ESPript 3.2. For phylogenetic analysis, homologous sequences were aligned using MUSCLE, and the tree was constructed in MEGA 12 using the Neighbor-Joining method (46). Evolutionary distances were calculated using the Poisson correction model, and bootstrap support values were estimated from 1000 replicates.

### Cryo-EM data collection and processing

Purified *V. stellata* PSI–VCPI complex (4.0 μL, corresponding to 1.5 mg Chl mL^-1^) was applied to glow-discharged Quantifoil Au R2/1 200-mesh grids and vitrified using a Vitrobot Mark IV (ThermoFisher Scientific) operated at 4°C and >90% relative humidity with 2 s blot time and −1 blot force. Cryo-EM data were collected on a 300 kV Titan Krios G4 transmission electron microscope (Thermo Fisher Scientific) equipped with a Falcon 4i BioQuantum direct electron detector. Movies were recorded at a nominal magnification of ×130,000, corresponding to a calibrated pixel size of 0.9275 Å. A total of 8,217 movie stacks were collected with a defocus range of −1.0 to −2.0 μm, a total electron dose of 60 e⁻Å^-2^, and a 20 eV energy slit.

Image processing was performed primarily in cryoSPARC v4.7.1 (47). Movie stacks were motion-corrected, dose-weighted, and then the contrast transfer function (CTF) parameters were estimated using Patch CTF (48, 49). Following particle picking, extracted particles were subjected to two rounds of reference-free 2D classification to remove false-positive particles and poorly defined classes. A total of 239,452 particles were selected for ab initio reconstruction and subsequent 3D classification. After removal of heterogeneous particles, the final dataset comprising 151,622 particles was subjected to homogeneous refinement. To further improve the map quality, non-uniform refinement, map sharpening, global (per-group) CTF refinement, local (per-particle) CTF refinement, particle subtraction, and masked local refinement were performed sequentially. The final reconstruction reached an overall resolution of 2.44 Å, as estimated by the gold-standard Fourier shell correlation (FSC) criterion at 0.143. The resolutions of the local refined region were 2.97 Å, 2.83 Å and 2.68 Å.

### Model building and refinement

The 2.44 Å resolution cryo-EM map of *V. stellata* PSI–VCPI was used for atomic model building. The *I. galbana* PSI–iFCPI structure (PDB ID: 8Z11) was first fitted into the cryo-EM density map using UCSF Chimera 1.12 (13, 50). The amino acid sequences of the PSI core subunits were then manually corrected in Coot according to the sequences obtained from the *V. stellata* transcriptome (51, 52). The three VCPI subunits were built using the corresponding antenna subunits of *I. galbana* PSI–iFCPI as starting models and were adjusted based on the agreement between the cryo-EM densities and the transcriptome-derived sequences. Only the N-terminal portion of FNR was modeled according to the corresponding transcriptome-derived sequence, whereas the remaining region was not built because of insufficient density. Chl *a*, *β*-carotene, violaxanthin, vaucheriaxanthin ester, phylloquinones, and Fe4S4 clusters were modeled into the densities. Chl *a* molecules and carotenoids were identified on the basis of density features. Geometrical restraints for pigments and cofactors were generated using the Grade Web Server. All residues, pigments, and cofactors were inspected and manually adjusted in Coot. The model was refined using Phenix real-space refinement, and the geometry of the final model was evaluated using Phenix (53, 54).

### Model construction and molecular dynamics simulations

Protein-membrane system was constructed using CHARMM-GUI and embedded in POPC lipid bilayers to obtain equilibrated protein–membrane complex structures for subsequent EET calculations (55), followed by solvation and neutralization. Before system construction, Ligand geometries were optimized at the DFT B3LYP/6-31G(d) level using Gaussian16 (56), and atomic charges were calculated using the restrained electrostatic potential method at the HF/6-31G(d) level with Gaussian16. Force-field parameters were then generated based on GAFF2 using the Antechamber module in AmberTools (57). The resulting ligand parameters were combined with the protein structures in tleap to generate the topology and coordinate files for subsequent system assembly. Proteins were described using the Amber ff14SB force field (58), POPC lipid molecules were described using the SLipids force field in an Amber-compatible format, and water molecules were modeled using the TIP3P water model.

The membrane orientation and transmembrane region of each protein complex were predicted using the PPM 2.0 method implemented in CHARMM-GUI. Protein structure was inserted into a prebuilt pure POPC bilayer. Unreasonable voids, gaps, or local packing defects at the protein–lipid interface were inspected and manually adjusted to ensure a physically reasonable membrane environment. The system is then solvated by adding a 10 Å water layer. Ions were added to neutralize the total charge and to adjust the ionic concentration to 0.15 M. Detailed information for each constructed system, including box dimensions, numbers of POPC lipids, water molecules, ions, and total atoms, is provided in Table S4.

Molecular dynamics simulations were performed using NAMD 3 under periodic boundary conditions (59), applying a two-stage equilibration protocol to relax the lipid environment and stabilize the protein-cofactor architecture. In the first stage, the backbone atoms of the protein and the retained pigments, cofactors, and other non-protein small molecules were kept fixed, whereas POPC lipids, water molecules, and ions were allowed to relax. This stage consisted of 50,000 steps of energy minimization, followed by 1 ns of NVT equilibration and 10 ns of NPT equilibration. In the second stage, the positional constraints were released, and the whole protein– membrane system was further equilibrated by 50,000 steps of energy minimization, 1 ns of NVT equilibration, and 30 ns of NPT equilibration. The detailed positional constraints applied at each equilibration step are summarized in Table S5. Five representative snapshots were selected from the last 10 ns of the final NPT trajectory at 2 ns intervals, after the protein backbone RMSD and key pigment–protein interactions had reached stable distributions (Fig. S14).

### QM/MM setup and structure optimization

To model the excited-state properties, a three-layer QM/MM approach was implemented using pDynamo2 integrated with BDF workflows (60–62). The system was partitioned into a QM region (the Chl macrocycle with the tail truncated at the first carbon atom C1 of the tail region and Mg-coordinating residues located within 3 Å of the central Mg atom), an active MM region (intact residues and molecules with at least one atom within 7 Å of the QM region), and a frozen MM region (intact residues or molecules located within 40 Å of the pigment molecule but outside the active MM region) to preserve the local electrostatic environment by being kept fixed (63).

Geometry optimizations were carried out sequentially—first relaxing the active MM region using standard force fields with a convergence threshold of 0.1 kJ mol^-1^ Å^-1^ and a maximum of 400 optimization steps, followed by optimizing the QM region using the semiempirical AM1 method under the electrostatic field of the MM point charges, with the root-mean-square gradient convergence threshold set to 0.01 kJ mol^-1^ Å^-1^ and the maximum number of optimization steps set to 2000. All QM/MM optimizations reached the predefined convergence criteria. The final optimized QM/MM structures were then used for subsequent TDDFT calculations, in which the pigment was calculated in the presence of the MM point-charge environment.

### Excitation energy transfer calculations

The optimized QM/MM structures were used for excitation energy transfer calculations. For each pigment, the first excited-state energy was evaluated using time-dependent density functional theory (64). The CAM-B3LYP functional was employed together with the 6-31G* basis set (65), and the quantum chemical calculations were performed using Gaussian 16. The resulting excitation energy was used as the corresponding pigment site energy. Pigment transition charges were derived using Multiwfn (66) for subsequent electronic coupling calculations. Pairwise electronic couplings between pigments were estimated using the transition charge from electrostatic potential method (67) and calculated with customized Python scripts. For each of the five representative structures, the site energies, transition charges, and pairwise electronic couplings were calculated independently. The final site energies and electronic coupling matrices used for subsequent EET analyses were obtained by averaging the corresponding values over the five structures, and the conformational variations were evaluated using their standard deviations.

Based on the calculated site energies and electronic couplings, pigment-to-pigment EET time constants were first evaluated using Förster resonance energy transfer theory (35). These pairwise transfer time constants were used to describe the local energy-transfer network among neighboring pigments.

Furthermore, pigment clusters or functional domains were defined according to the structural organization of the protein complex (36). For each cluster, the pigment site energies and intracluster electronic couplings were used to construct the excitonic Hamiltonian, from which cluster exciton states were obtained. Generalized Förster theory was then applied to calculate cluster-level energy transfer time constants between donor and acceptor domains, using the exciton-state energies, interdomain electronic couplings, and spectral overlap factors. The overall computational protocol closely followed those used in our previous studies (11, 13, 68).

## Data availability

The cryo-EM map and atomic coordinates generated in this study have been deposited in the Protein Data Bank and Electron Microscopy Data Bank under the accession numbers 43QU and EMD-82074, respectively.

## Acknowledgments

We thank Qiu-Yao Jiang (Cryo-Electron Microscopy Platform of Medical Science and Technology Innovation Center of Shandong First Medical University), Xiao-Ju Li (State Key Laboratory of Microbial Technology, Shandong University, Qingdao, China) and Dian-Li Zhao (Laoshan Laboratory, Qingdao, Shandong, China) for their contributions in cryo-EM data collection. Numerical computations were performed on Hefei advanced computing center. This work was supported by National Key R&D Program of China (2023YFA0914600 to LSZ and LNL), Shandong Province Science Fund for Excellent Young Scholars (ZR2024YQ027 to LSZ), National Natural Science Foundation of China (32570122 to LSZ, 32330001 to YZZ, W2441012 to YZZ, 32070109 to LNL, 21873034 to JG), National Key R&D Program of China (2021YFA0909600 to LNL), the Taishan Scholars Program of Shandong Province, China (tsqn202306092 to YZZ), Natural Science Foundation of Shandong Province (ZR2024QC042 to KL), the SKLMT Frontiers and Challenges Project (SKLMTFCP-2023-06 to YZZ, Qilu Youth Scholar Startup Funding of SDU (to LSZ), the Royal Society (URF\R\180030 to LNL), Leverhulme Trust (RPG-2021-286 to LNL), Biotechnology and Biological Sciences Research Council (BBSRC) (BB/Y01135X/1, UKRI4524, and BB/W001012/1 to LNL), Fundamental Research Funds for the Central Universities (2662024XXPY003).

## Author Contributions

Y.-Z.Z. and L.-S.Z. conceived the project. Y.L., L.-S.Z., and X.-M.S. performed sample preparation and measurements. L.-D.H. kindly provided the algal strain and instructed the algal culture and cell-disruption experiments. K.L. collected and processed the cryo-EM data. Y.L. and L.-S.Z. built the structural model and refined the structure. Q.W. and J.G. performed computational simulations for excitation energy transfer. L.-S.Z., K.L., Y.L., F. Z., X.-X.Q., and H.-J.W. analyzed the data. L.-S.Z., Y.L., Q.W., L.-N.L., and Y.-Z.Z. wrote the manuscript, with contributions from all other authors.

## References

1. P. Sétif, “Ferredoxin and flavodoxin reduction by photosystem I” in Biochim Biophys Acta. (Netherlands, 2001), vol. 1507, pp. 161–179.

2. P. Mulo, “Chloroplast-targeted ferredoxin-NADP(+) oxidoreductase (FNR): structure, function and location” in Biochim Biophys Acta. (Netherlands, 2011), vol. 1807, pp. 927 –934.

3. X. Pi et al., “Unique organization of photosystem I-light-harvesting supercomplex revealed by cryo-EM from a red alga” in Proc Natl Acad Sci U S A. (United States, 2018), vol. 115, pp. 4423 –4428.

4. M. Antoshvili, I. Caspy, M. Hippler, N. Nelson, “Structure and function of photosystem I in Cyanidioschyzon merolae” in Photosynth Res. (Netherlands, 2019), vol. 139, pp. 499 –508.

5. X. You et al., “In situ structure of the red algal phycobilisome-PSII-PSI-LHC megacomplex” in Nature. (England, 2023), vol. 616, pp. 199–206.

6 K. Kato et al., “The structure of PSI-LHCI from Cyanidium caldarium provides evolutionary insights into conservation and diversity of red-lineage LHCs” in Proc Natl Acad Sci U SA. (United States, 2024), vol. 121, pp. e2319658121.

7. K. Kato, Structure of a photosystem I supercomplex from Galdieria sulphuraria close to an ancestral red alga. Science Advances 10.1126/sciadv.adv7488 (2025).

8. L. S. Zhao et al., “Structural basis and evolution of the photosystem I-light-harvesting supercomplex of cryptophyte algae” in Plant Cell. (England, 2023), vol. 35, pp. 2449 –2463.

9. S. Zhang et al., “Growth phase-dependent reorganization of cryptophyte photosystem I antennae” in Commun Biol. (England, 2024), vol. 7, pp. 560.

10. W. Zhang et al., Structural analysis of PSI-ACPI and PSII-ACPII supercomplexes from a cryptophyte alga Rhodomonas sp. NIES-2332. Front Plant Sci 16, 1716939 (2025).

11. X. M. Sun et al., “Architecture and energy transfer of coccolithophore photosystem I with a huge light-harvesting antenna system” in Sci Adv. (United States, 2025), vol. 11, pp. eaea4965.

12. L. Shen et al., Structure and function of a huge photosystem I-fucoxanthin chlorophyll supercomplex from a coccolithophore. Science 389, eadv2132 (2025).

13. F. Y. He et al., “Structural insights into the assembly and energy transfer of haptophyte photosystem I -light-harvesting supercomplex” in Proc Natl Acad Sci U S A. (United States, 2024), vol. 121, pp. e2413678121.

14. C. Xu et al., “Structural basis for energy transfer in a huge diatom PSI-FCPI supercomplex” in Nat Commun. (England, 2020), vol. 11, pp. 5081.

15. R. Nagao et al., “Structural basis for assembly and function of a diatom photosystem I -light-harvesting supercomplex” in Nat Commun. (England, 2020), vol. 11, pp. 2481.

16. K. Kato et al., “Structural basis for molecular assembly of fucoxanthin chlorophyll a/c-binding proteins in a diatom photosystem I supercomplex” in Elife. (England, 2024), vol. 13.

17. Y. Feng et al., Structures of PSI-FCPI from Thalassiosira pseudonana grown under high light provide evidence for convergent evolution and light-adaptive strategies in diatom FCPIs. J Integr Plant Biol 67, 949–966 (2025).

18. R. Shao et al., Architecture of photosystem I-light-harvesting complex from the eukaryotic filamentous yellow-green alga Tribonema minus. J Integr Plant Biol 67, 3014–3031 (2025).

19. J. D. Weissman et al., “Structure of giant kelp Photosystem I-FCP uncovers drivers of antenna evolution across the red lineage” in Nat Commun. (England, 2026), 10.1038/s41467-026-73499-x.

20. L. S. Zhao et al., “Architecture of symbiotic dinoflagellate photosystem I-light-harvesting supercomplex in Symbiodinium” in Nat Commun. (England, 2024), vol. 15, pp. 2392.

21. X. Li et al., “Structures and organizations of PSI-AcpPCI supercomplexes from red tidal and coral symbiotic photosynthetic dinoflagellates” in Proc Natl Acad Sci U S A. (United States, 2024), vol. 121, pp. e2315476121.

22. V. E. J. Jassey et al., Contribution of soil algae to the global carbon cycle. New Phytol 234, 64–76 (2022).

23. S. Aigner et al., Adaptation to Aquatic and Terrestrial Environments in Chlorella vulgaris (Chlorophyta). Front Microbiol 11, 585836 (2020).

24. I. Caspy et al., “Cryo-EM photosystem I structure reveals adaptation mechanisms to extreme high light in Chlorella ohadii” in Nat Plants. (England, 2021), vol. 7, pp. 1314 –1322.

25. K. X. Terpis et al., Multiple plastid losses within photosynthetic stramenopiles revealed by comprehensive phylogenomics. Curr Biol 35, 483–499 e488 (2025).

26. A. Cho, G. Lax, P. J. Keeling, Phylogenomic analyses of ochrophytes (stramenopiles) with an emphasis on neglected lineages. Molecular Phylogenetics and Evolution 198 (2024).

27. M. Eliáš et al., “Eustigmatophyceae” in Handbook of the Protists. (2017), 10.1007/978-3-319-32669-6_39-1 chap. Chapter 39-1, pp. 1–39.

28. S. Takaichi, “Carotenoids in algae: distributions, biosyntheses and functions” in Mar Drugs. (Switzerland, 2011), vol. 9, pp. 1101–1118.

29. R. Amaral, J. S. S. de Melo, L. M. A. dos Santos, Pigments from Eustigmatophyceae: an interesting class of microalgae for carotenoid production. Journal of Applied Phycology 33, 371–384 (2021).

30. A. Sukenik, A. Livne, K. E. Apt, A. R. Grossman, Characterization of a Gene Encoding the Light-Harvesting Violaxanthin-Chlorophyll Protein of Nannochloropsis Sp. (Eustigmatophyceae). J Phycol 36, 563–570 (2000).

31. X. Pan et al., Structural basis for energy transfer and electron transport in the photosystem I-ferredoxin-NADP+ reductase supercomplex from Nannochloropsis oceanica. 10.21203/rs.3.rs-8964687/v1 (2026).

32. B. Gao et al., Characterization of cell structural change, growth, lipid accumulation, and pigment profile of a novel oleaginous microalga, Vischeria stellata (Eustigmatophyceae), culturedwith differentinitialnitratesupplies. Journal of Applied Phycology 28, 821–830 (2015).

33. R. Morales, G. Kachalova, F. Vellieux, M. H. Charon, M. Frey, “Crystallographic studies of the interaction between the ferredoxin-NADP+ reductase and ferredoxin from the cyanobacterium Anabaena: looking for the elusive ferredoxin molecule” in Acta Crystallogr D Biol Crystallogr. (United States, 2000), vol. 56, pp. 1408 – 1412.

34. G. Kurisu et al., Structure of the electron transfer complex between ferredoxin and ferredoxin-NADP(+) reductase. Nat Struct Biol 8, 117–121 (2001).

35. M. Sener et al., Forster energy transfer theory as reflected in the structures of photosynthetic light-harvesting systems. Chemphyschem 12, 518–531 (2011).

36. J. Hsin et al., Energy Transfer Dynamics in an RC-LH1-PufX Tubular Photosynthetic Membrane. New J Phys 12 (2010).

37. R. Derelle, P. Lopez-Garcia, H. Timpano, D. Moreira, “A Phylogenomic Framework to Study the Diversity and Evolution of Stramenopiles (=Heterokonts)” in Mol Biol Evol. (United States, 2016), vol. 33, pp. 2890 –2898.

38. J. I. Kim et al., “Evolutionary Dynamics of Cryptophyte Plastid Genomes” in Genome Biol Evol. (England, 2017), vol. 9, pp. 1859–1872.

39. J. W. Stiller et al., “The evolution of photosynthesis in chromist algae through serial endosymbioses” in Nat Commun. (England, 2014), vol. 5, pp. 5764.

40. M. Kumazawa, K. Ifuku, “Unraveling the evolutionary trajectory of LHCI in red-lineage algae: Conservation, diversification, and neolocalization” in iScience. (United States, 2024), vol. 27, pp. 110897.

41. X. Pan et al., “Structure of the maize photosystem I supercomplex with light-harvesting complexes I and II” in Science. (United States, 2018), vol. 360, pp. 1109–1113.

42. X. Qin et al., “Structureof a greenalgal photosystem I in complex with a large number of light-harvesting complex I subunits” in Nat Plants. (England, 2019), vol. 5, pp. 263 –272.

43. X. Qin, M. Suga, T. Kuang, J. R. Shen, “Photosynthesis. Structural basis for energy transfer pathways in the plant PSI-LHCI supercomplex” in Science. (United States, 2015), vol. 348, pp. 989 –995.

44. X. Su et al., “Antenna arrangement and energy transfer pathways of a green algal photosystem-I-LHCI supercomplex” in Nat Plants. (England, 2019), vol. 5, pp. 273 –281.

45. M. Stoyneva-Gärtner et al., Carotenoids in five aeroterrestrial strains fromVischeria/Eustigmatosgroup: updating the pigment pattern of Eustigmatophyceae. Biotechnology & Biotechnological Equipment 33, 250–267 (2019).

46. S. Kumar, et al., “MEGA12: Molecular Evolutionary Genetic Analysis Version 12 for Adaptive and Green Computing” in Mol Biol Evol. (United States, 2024), vol. 41.

47. A. Punjani, J. L. Rubinstein, D. J. Fleet, M. A. Brubaker, “cryoSPARC: algorithms for rapid unsupervised cryo-EM structure determination” in Nat Methods. (United States, 2017), vol. 14, pp. 290 –296.

48. S. Q. Zheng et al., “MotionCor2: anisotropic correction of beam-induced motion for improved cryo-electron microscopy” in Nat Methods. (United States, 2017), vol. 14, pp. 331 –332.

49. A. Rohou, N. Grigorieff, CTFFIND4: Fast and accurate defocus estimation from electron micrographs. J Struct Biol 192, 216–221 (2015).

50. E. F. Pettersen et al., UCSF Chimera--a visualization system for exploratory research and analysis. J Comput Chem 25, 1605–1612 (2004).

51. A. Casañal, B. Lohkamp, P. Emsley, “Current developments in Coot for macromolecular model building of Electron Cryo-microscopy and Crystallographic Data” in Protein Sci. (United States, 2020), vol. 29, pp. 1069–1078.

52. P. Emsley, B. Lohkamp, W. G. Scott, K. Cowtan, “Features and development of Coot” in Acta Crystallogr D Biol Crystallogr. (United States, 2010), vol. 66, pp. 486–501.

53. P. D. Adams et al., “PHENIX: a comprehensive Python-based system for macromolecular structure solution” in Acta Crystallogr D Biol Crystallogr. (United States, 2010), vol. 66, pp. 213 –221.

54. P. D. Adams et al., “The Phenix software for automated determination of macromolecular structures” in Methods. (United States, 2011), vol. 55, pp. 94–106.

55. S. Feng, S. Park, Y. K. Choi, W. Im, CHARMM-GUI Membrane Builder: Past, Current, and Future Developments and Applications. J Chem Theory Comput 19, 2161–2185 (2023).

56. M. J. Frisch et al. (2016) Gaussian 16, Revision C.01. (Gaussian, Inc., Wallingford, CT).

57. D. A. Case et al., AmberTools. Journal of Chemical Information and Modeling 63, 6183–6191 (2023).

58. J. A. Maier et al., ff14SB: Improving the Accuracy of Protein Side Chain and Backbone Parameters from ff99SB. Journal of Chemical Theory and Computation 11, 3696–3713 (2015).

59. J. C. Phillips et al., “Scalable molecular dynamics on CPU and GPU architectures with NAMD” in J Chem Phys. (United States, 2020), vol. 153, pp. 044130.

60. M. J. Field, M. Albe, C. l. Bret, F. Proust-De Martin, A. Thomas, The dynamo library for molecular simulations using hybrid quantum mechanical and molecular mechanical potentials. Journal of Computational Chemistry 21, 1088–1100 (2000).

61. M. J. Field, The pDynamo Program for Molecular Simulations using Hybrid Quantum Chemical and Molecular Mechanical Potentials. Journal of Chemical Theory and Computation 4, 1151–1161 (2008).

62. Y. Zhang et al., BDF: A relativistic electronic structure program package. The Journal of Chemical Physics 152 (2020).

63. J.-P. Guo et al., Multi-scale simulations of excited-state energy transfer pathways in the C2S2-type PSII–LHCII supercomplex of spinach. Physical Chemistry Chemical Physics 28, 5479–5490 (2026).

64. E. Runge, E. K. U. Gross, Density-Functional Theory for Time-Dependent Systems. Physical Review Letters 52, 997–1000 (1984).

65. T. Yanai, D. P. Tew, N. C. Handy, A new hybrid exchange–correlation functional using the Coulomb-attenuating method (CAM-B3LYP). Chemical Physics Letters 393, 51–57 (2004).

66. T. Lu, F. Chen, Multiwfn: Amultifunctional wavefunction analyzer. Journal of Computational Chemistry 33, 580– 592 (2012).

67. M. E. Madjet, A. Abdurahman, T. Renger, Intermolecular Coulomb Couplings from Ab Initio Electrostatic Potentials: Application to Optical Transitions of Strongly Coupled Pigments in Photosynthetic Antennae and Reaction Centers. The Journal of Physical Chemistry B 110, 17268–17281 (2006).

68. K. Li et al., “Structure and energy transfer of a far-red-absorbing euglenophyte PSI-LhcE-LhcbM supercomplex” in Nat Commun. (England, 2026), vol. 17.

