## Supplemental Figures and Tables for "Structure and energy transfer of a minimal PSI–LHCI supercomplex with FNR binding from a terrestrial eustigmatophyte"

**This PDF file includes:**

Figs. S1 to S14
Tables S1 to S5

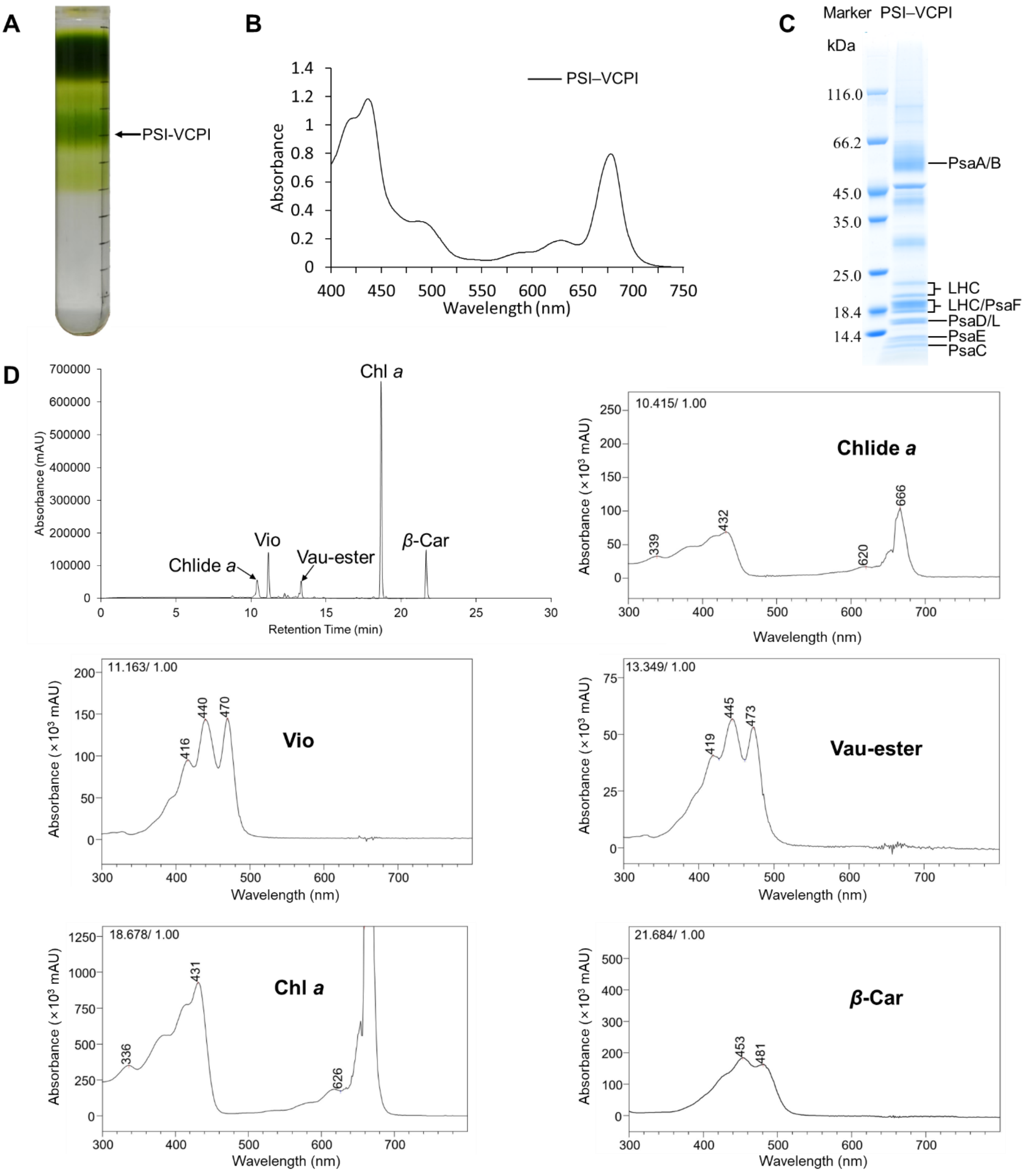

**Fig. S1. Preparation and characterization of PSI–VCPI from *V. stellata*. (A)** Isolation of the PSI–VCPI supercomplex by ultracentrifugation using a sucrose density gradient. **(B)** Room-temperature absorption spectrum of the purified PSI–VCPI supercomplex. **(C)** SDS-PAGE analysis of the purified PSI–VCPI supercomplex. The protein composition of the bands was assigned based on mass spectrometry analysis. **(D)** Pigment analysis of PSI–VCPI by high-performance liquid chromatography (HPLC), recorded at 440 nm. Based on the characteristic absorption spectra of the peak fractions, the major pigment peaks were identified as chlorophyllide *a* (Chlide *a*), violaxanthin (Vio), vaucheriaxanthin ester (Vau-ester), chlorophyll *a* (Chl *a*), and *β*-carotene (*β*-Car). Absorption spectra of these peaks are shown.

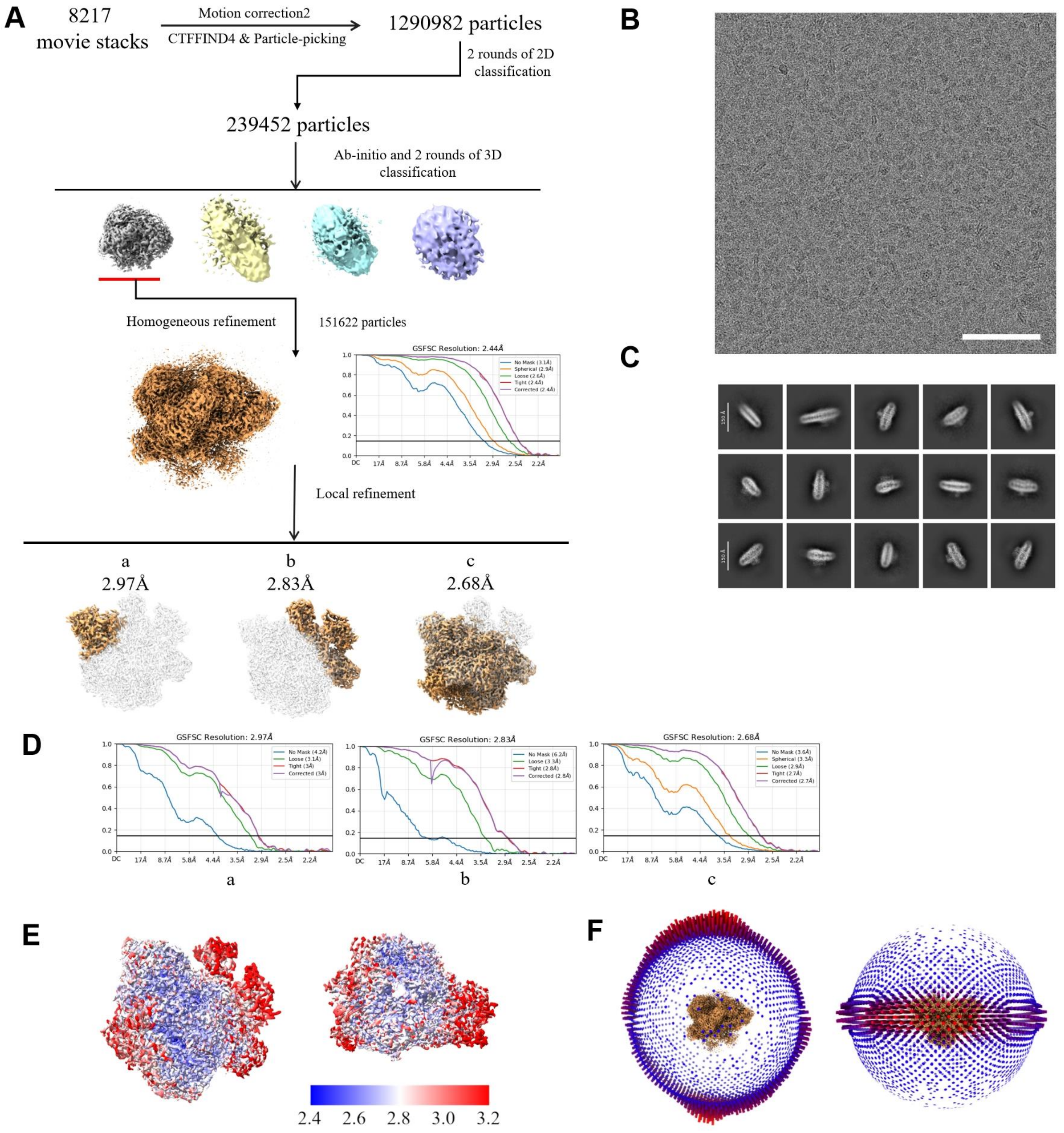

**Fig. S2. Cryo-EM data processing for the eustigmatophyte VsPSI–VCPI supercomplex. (A)** Schematic flowchart for the cryo-EM data processing. **(B)** A representative cryo-EM micrograph of the eustigmatophyte PSI–VCPI supercomplex. Scale bar, 100 nm. **(C)** Representative 2D classes of the eustigmatophyte PSI–VCPI supercomplex. The box size is 360 pixel approximately equal to 340 Å. **(D)** The gold standard Fourier shell correlation (FSC) curves for estimation of the resolution of the local density map with criterion of 0.143. **(E)** Local resolution distributions of the cryo-EM map estimated by ResMap. **(F)** Angular distribution of particles used for reconstruction of the final density map.

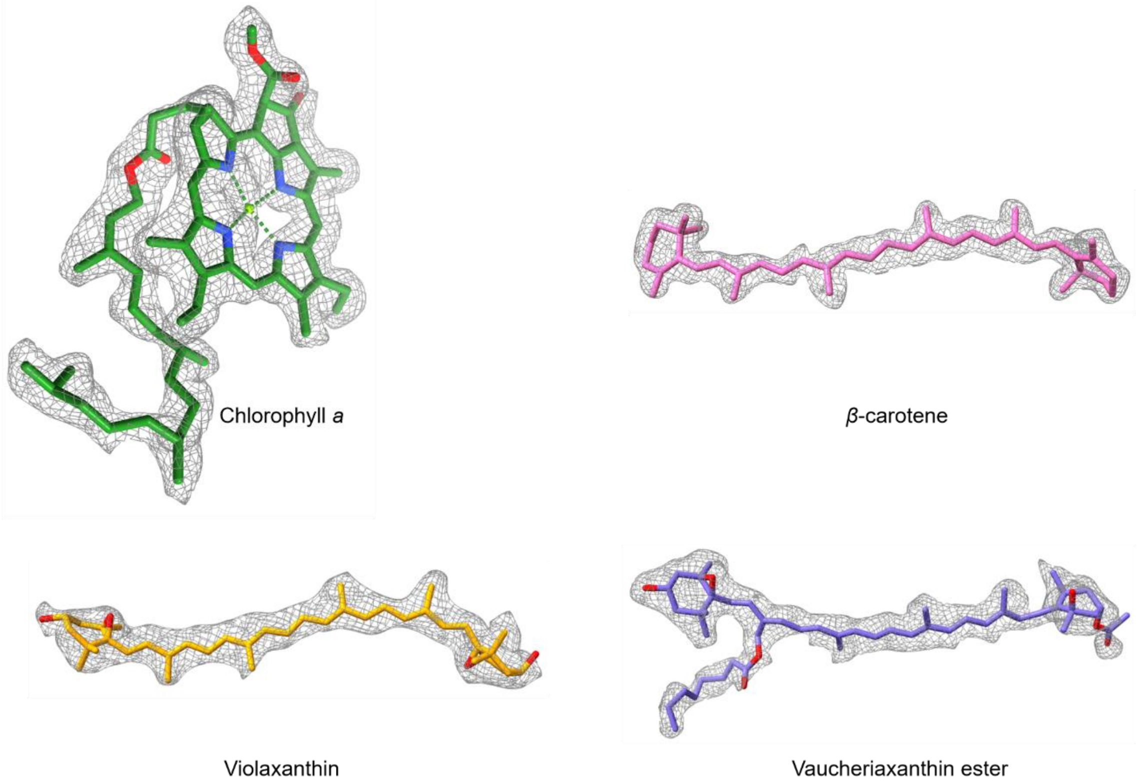

**Fig. S3. Cryo-EM density maps and structures of pigment molecules in the eustigmatophyte VsPSI–VCPI supercomplex.**

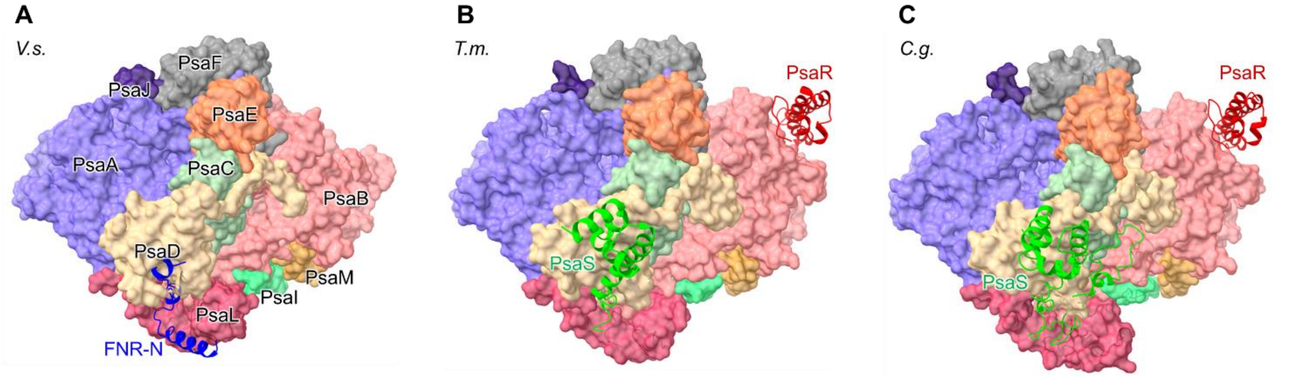

**Fig. S4. Comparison of PSI core structures from *V. stellata*, *T. minus*, and *C. gracilis*. (A–C)** Stromal-side views of the PSI core structures from *V. stellata* (*V.s.*) (A), *Tribonema minus* (*T.m.*) (B), and *Chaetoceros gracilis* (*C.g.*) (C). FNR-N in *V. stellata* is shown as blue cartoons. PsaS and PsaR in *T. minus* and *C. gracilis* are shown as green and red cartoons, respectively.

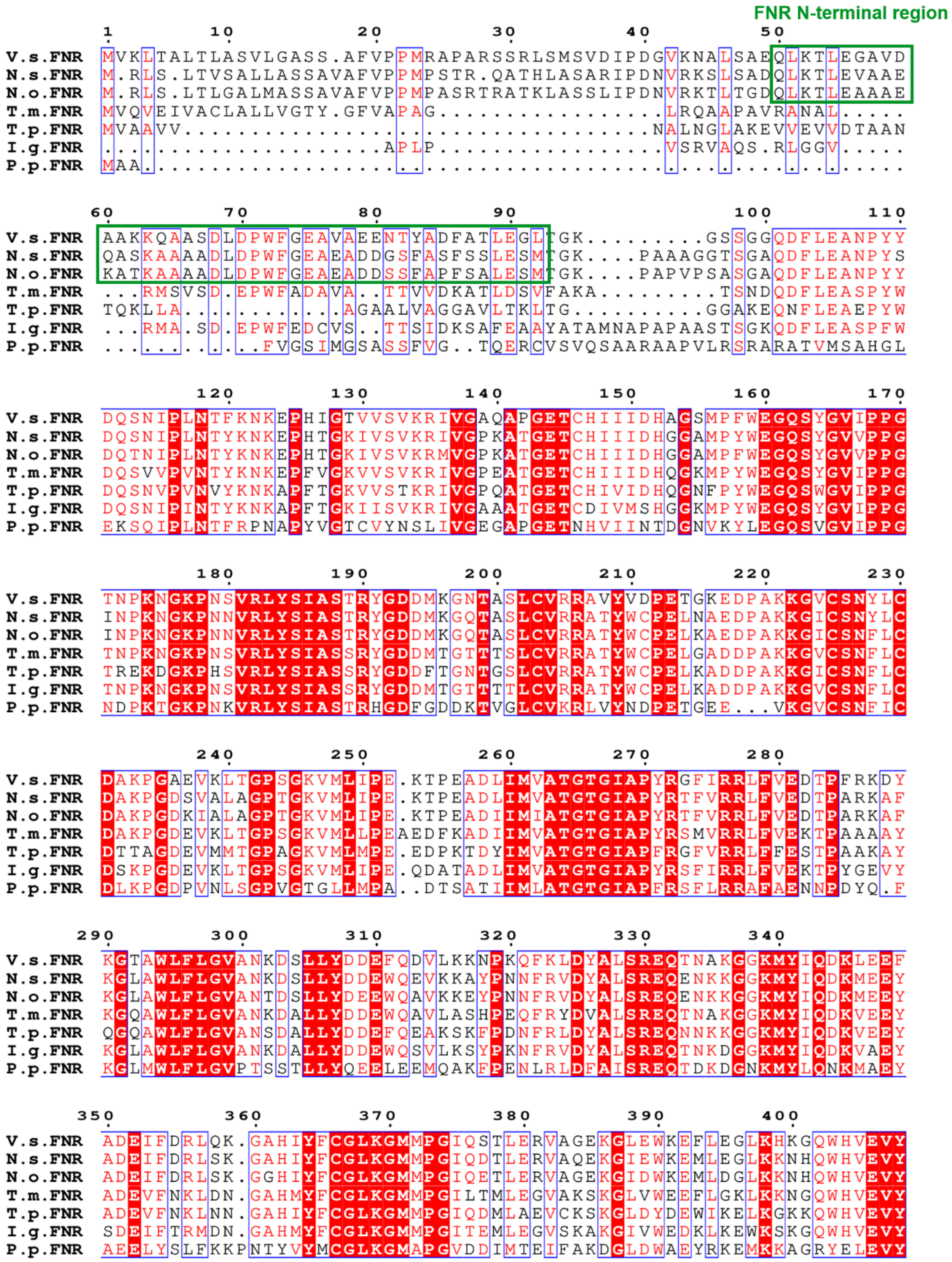

**Fig. S5. Sequence alignment of FNR from *V. stellata* and other red-lineage algae.** The sequence alignment of FNR in *Vischeria stellata* (*V.s.*), *Nannochloropsis salina* (*N.s.*), *Nannochloropsis oceanica* (*N.o.*), *Tribonema minus* (*T.m.*), *Thalassiosira pseudonana* (*T.p.*), *Isochrysis galbana* (*I.g.*) and *Porphyridium purpureum* (*P.p.*). The green boxes indicate the N-terminal region of FNR that is conserved in eustigmatophytes and corresponds to the FNR-N region modeled in *V. stellata* PSI–VCPI structure. Identical and similar residues are highlighted in red and boxed in blue, respectively. Dots indicate gaps introduced for sequence alignment.

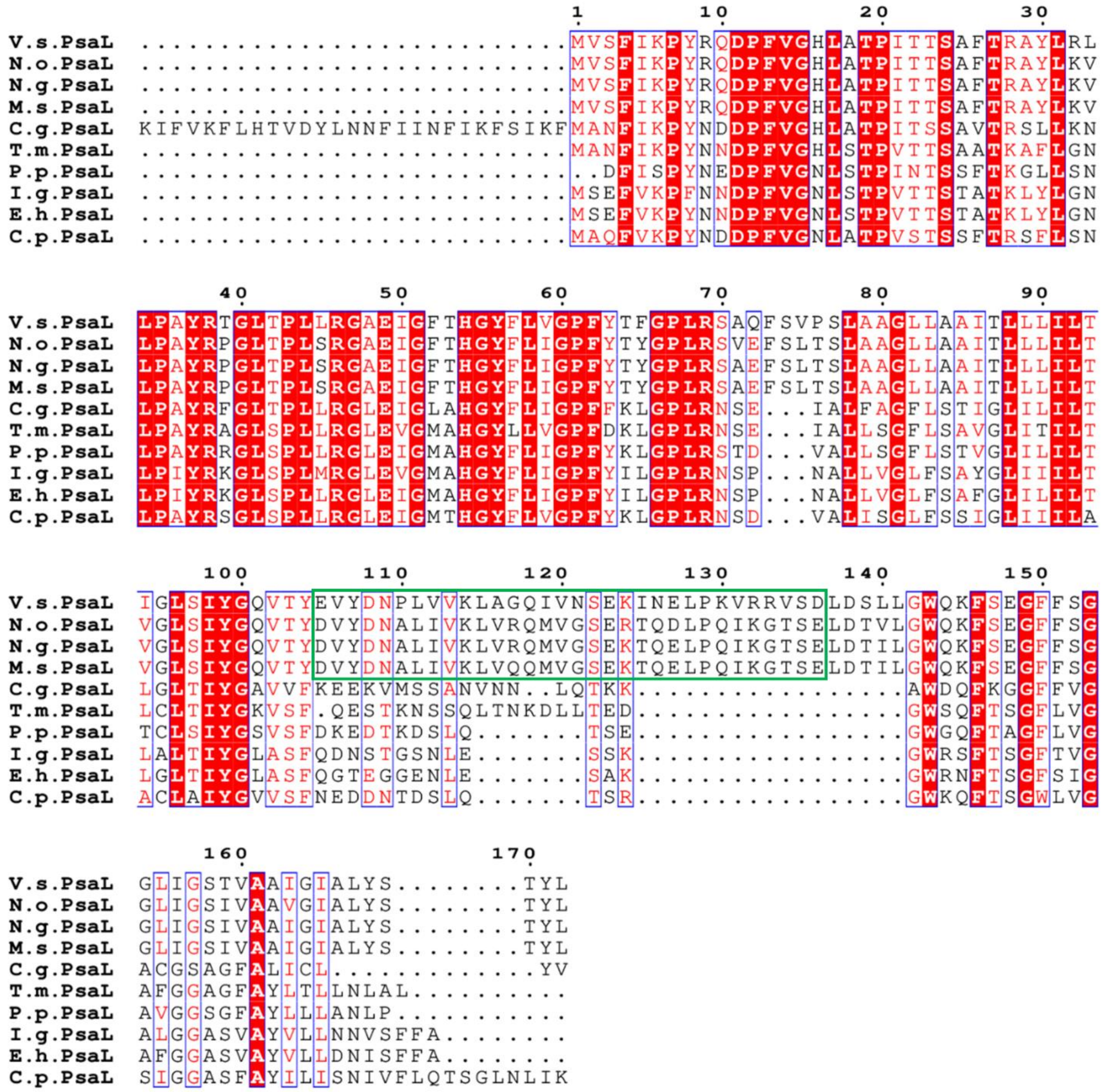

**Fig. S6. Sequence alignment of PsaL from *V. stellata* and other red-lineage algae.** The sequence alignment of PsaL in *Vischeria stellata* (*V.s.*), *Nannochloropsis oceanica* (*N.o.*), *Nannochloropsis gaditana* (*N.g.*), *Microchloropsis salina* (*M.s.*), *Chaetoceros gracilis* (*C.g.*), *Tribonema minus* (*T.m.*), *Porphyridium purpureum* (*P.p.*), *Isochrysis galbana* (*I.g.*), *Emiliania huxleyi* (*E.h.*) and *Chroomonas placoidea* (*C.p.*). The green box indicates the specific extended loop of PsaL conserved in eustigmatophytes. Identical and similar residues are highlighted in red and boxed in blue, respectively. Dots indicate gaps introduced for sequence alignment.

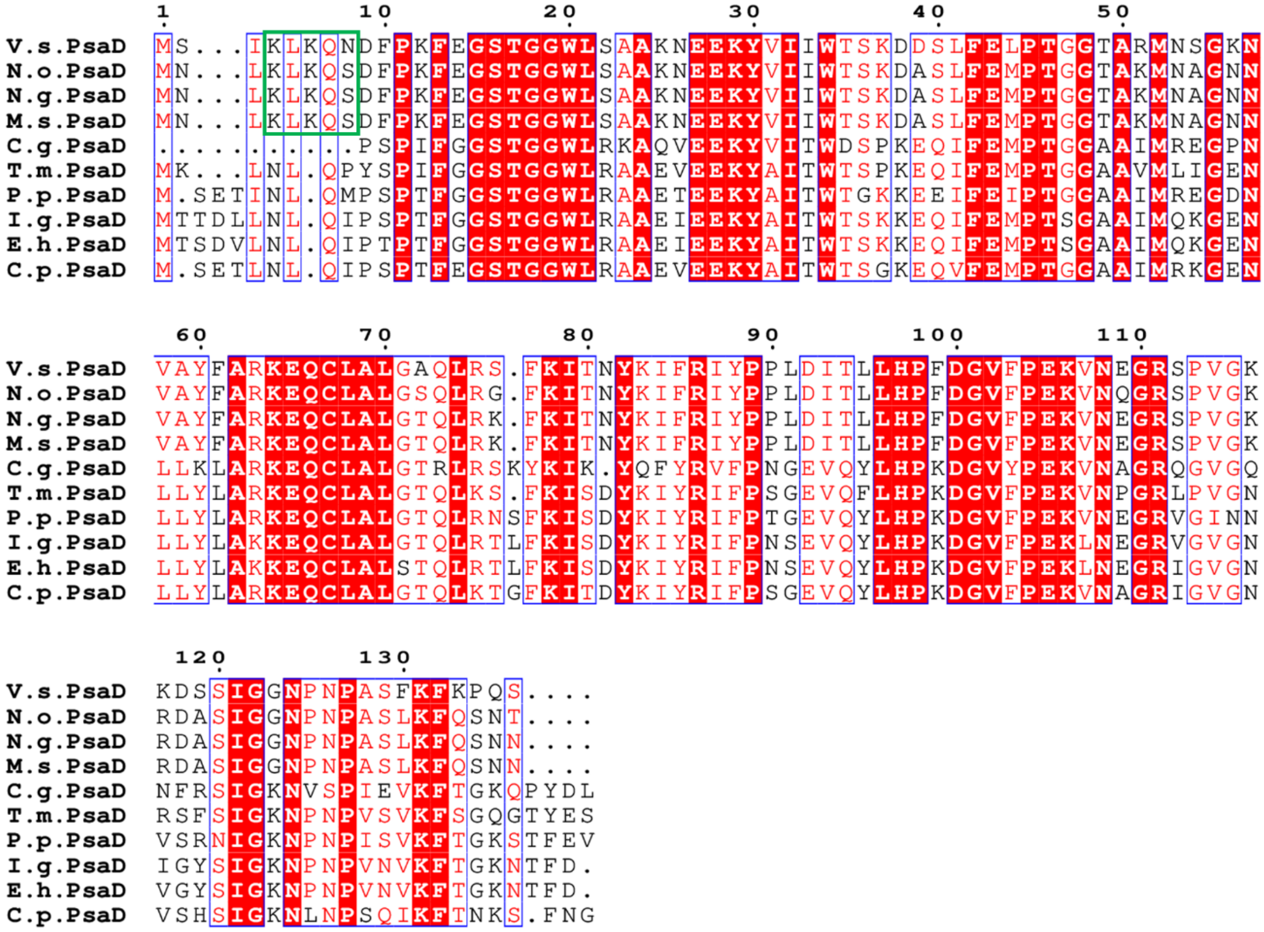

**Fig. S7. Sequence alignment of PsaD from *V. stellata* and other red-lineage algae.** The sequence alignment of PsaD in *Vischeria stellata* (*V.s.*), *Nannochloropsis oceanica* (*N.o.*), *Nannochloropsis gaditana* (*N.g.*), *Microchloropsis salina* (*M.s.*), *Chaetoceros gracilis* (*C.g.*), *Tribonema minus* (*T.m.*), *Porphyridium purpureum* (*P.p.*), *Isochrysis galbana* (*I.g.*), *Emiliania huxleyi* (*E.h.*) and *Chroomonas placoidea* (*C.p.*). The green box indicates the N-terminal region of PsaD that is conserved in eustigmatophytes and contributes to the interaction with FNR-N. Identical and similar residues are highlighted in red and boxed in blue, respectively. Dots indicate gaps introduced for sequence alignment.

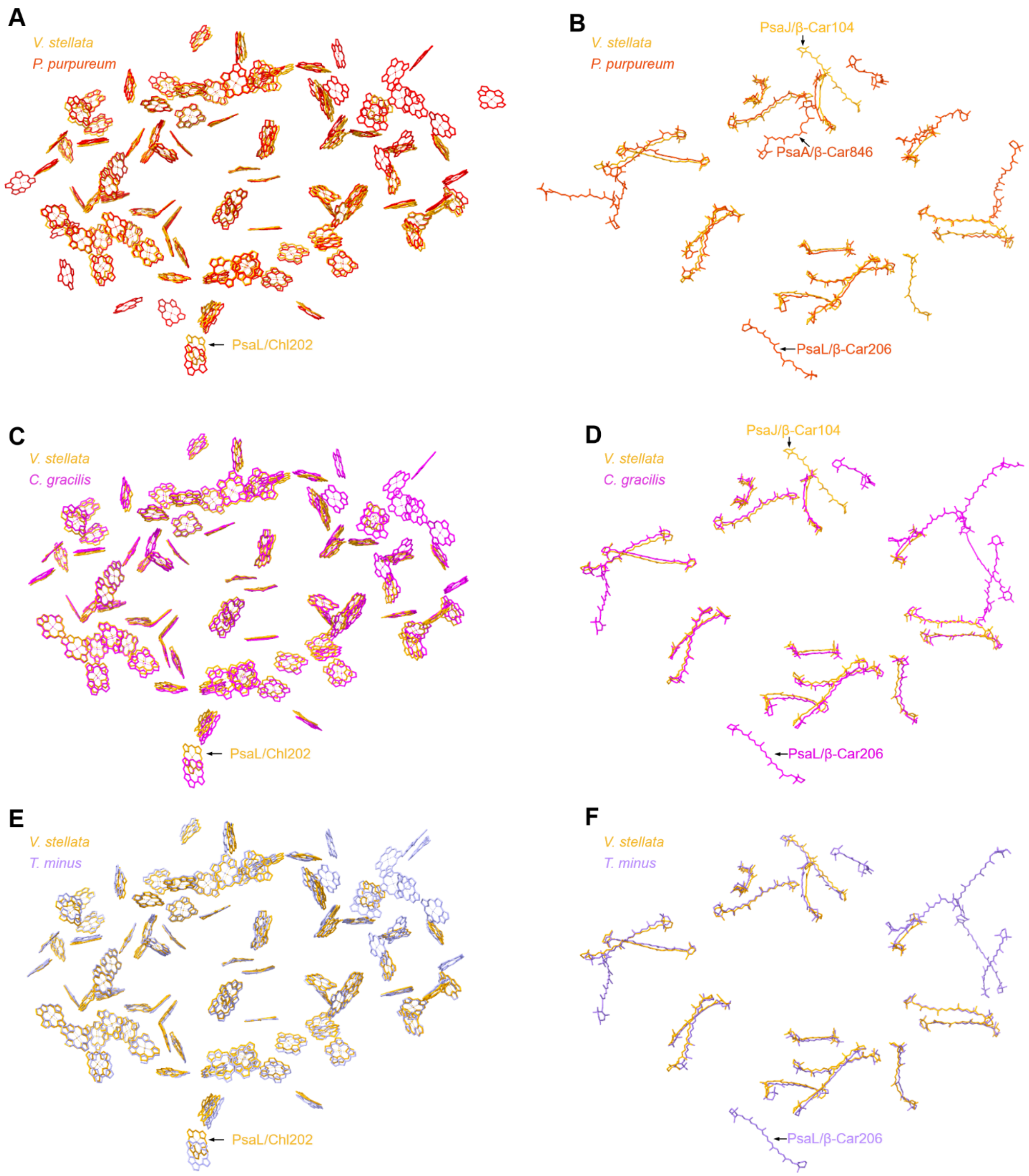

**Fig. S8. Comparison of pigment arrangements in *V. stellata* PSI core with those of red alga (PDB: 7Y5E), diatom (PDB: 6LY5) and xanthophyte (PDB: 9M4F).** Superpositions of the chlorophyll sites and carotenoid sites in *V. stellata* PSI core with those in red algae *Porphyridium purpureum* PSI core (red) **(A to B)**, diatom *Chaetoceros gracilis* PSI core (magenta) **(C to D)**, xanthophyte *Tribonema minus* PSI core (purple) **(E to F)**. Distinct sites in *V. stellata* PSI core are labeled. All panels are viewed from the stromal side.

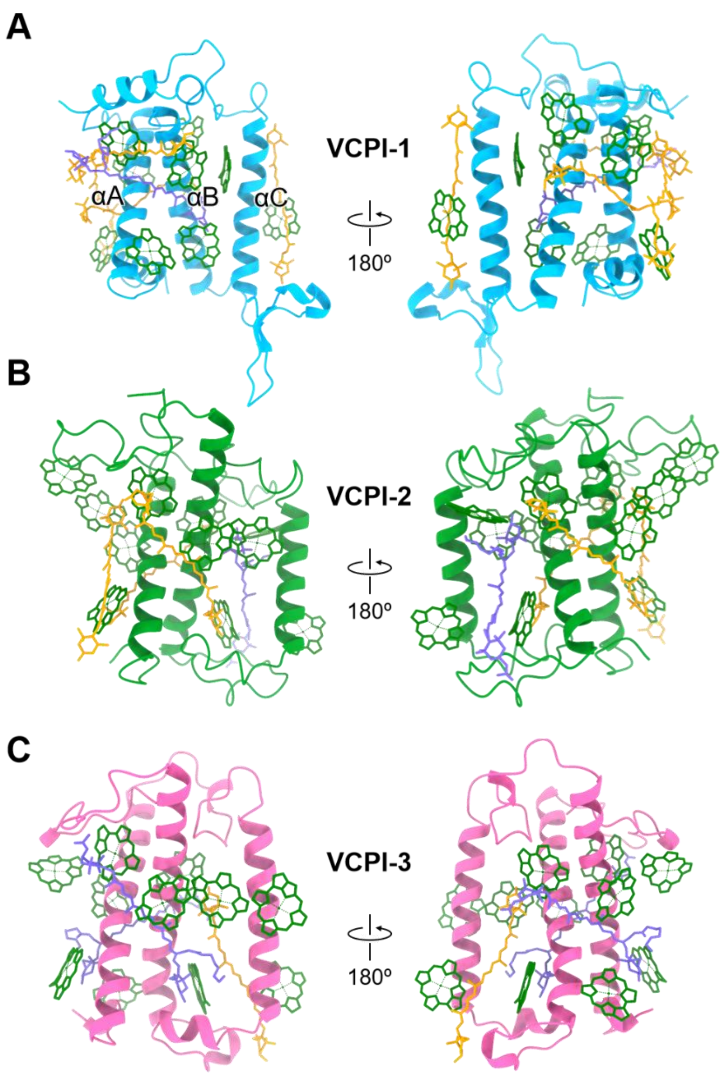

**Fig. S9. Structures of individual 3 VCPIs of eustigmatophyte PSI–VCPI. (A–C)** Structures of VCPI-1, VCPI-2, and VCPI-3, respectively. Each VCPI is shown in two opposite views related by a 180° rotation. Chlorophyll *a*, violaxanthin, and vaucheriaxanthin ester are colored green, yellow, and purple, respectively. The phytol chains of Chls are omitted.

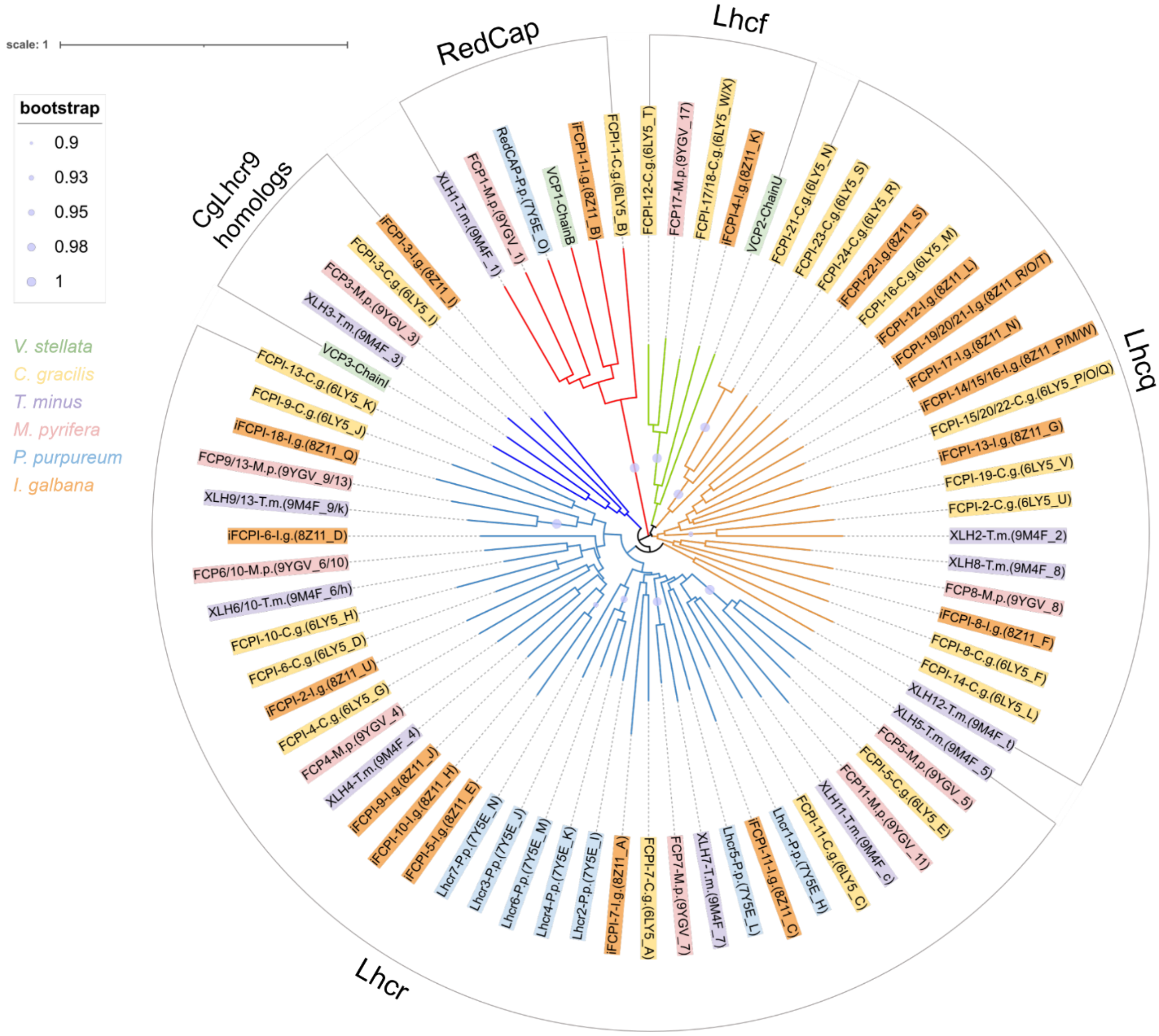

**Fig. S10. Phylogenetic tree of LHC antenna subunits in *Vischeria stellata* (*V.s.*), *Chaetoceros gracilis* (*C.g.*), *Tribonema minus* (*T.m.*), *Macrocystis pyrifera* (*M.p.*), *Porphyridium purpureum* (*P.p.*) and *Isochrysis galbana* (*I.g.*).** The Neighbor-Joining tree was constructed based on the amino acid sequences of LHC antenna subunits. The tree was built using 447 amino acid residues, and a bootstrap test (1000 replicates) was conducted. Chain ID of each VCPI is labeled. The PDB ID and Chain ID of FCPIs, XLHs, FCPs, Lhcrs and iFCPIs are indicated in the form of "(PDB ID_Chain ID)".

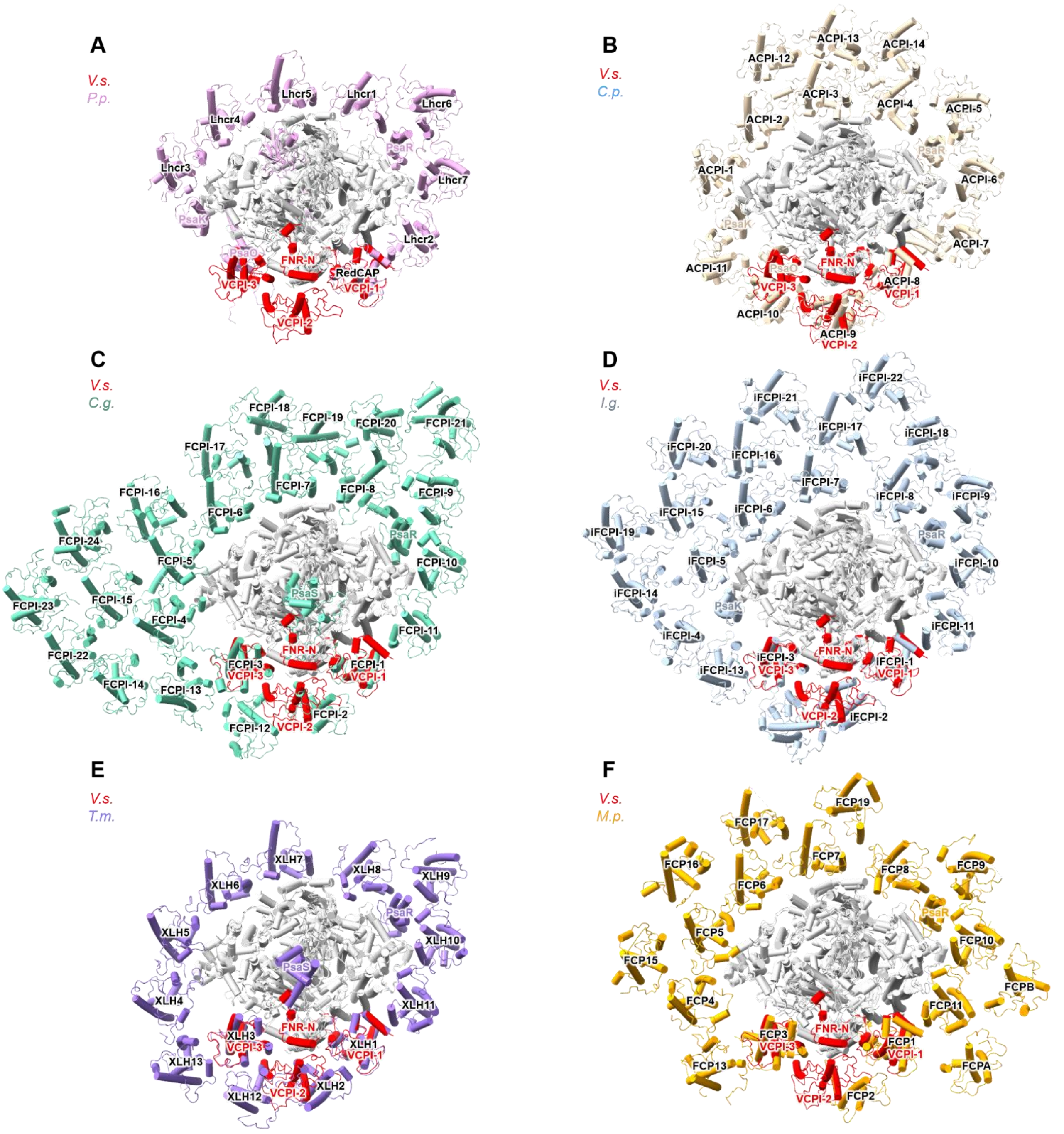

**Fig. S11. Comparison of LHCs arrangements in *Vischeria stellata* PSI–VCPI with those in *Porphyridium purpureum* (*P.p.*) PSI–LHCR (PDB: 7Y5E), *Chroomonas placoidea* (*C.p.*) PSI–ACPI (PDB: 7Y7B), *Chaetoceros gracilis* (*C.g.*) PSI–FCPI (PDB: 6LY5), *Isochrysis galbana* (*I.g.*) PSI–iFCPI (PDB: 8Z11), *Tribonema minus* (*T.m.*) PSI–XLH (PDB: 9M4F) and *Macrocystis pyrifera* (*M.p.*) PSI–FCP (PDB: 9YGV). (A)** Superposition of the *V. stellata* PSI–VCPI and *P. purpureum* PSI–LHCR structures. PsaK, PsaO, PsaR and Lhcrs in *P. purpureum* are labeled, and FNR-N and VCPIs in *V. stellata* are labeled. **(B)** Superposition of the *V. stellata* PSI–VCPI and *C. placoidea* PSI–ACPI structures. PsaK, PsaO, PsaR and ACPIs in *C. placoidea* are labeled. **(C)** Superposition of the *V. stellata* PSI–VCPI and *C. gracilis* PSI–FCPI structures. PsaS, PsaR and FCPIs in *C. gracilis* are labeled. **(D)** Superposition of the *V. stellata* PSI–VCPI and *I. galbana* PSI–iFCPI structures. PsaK, PsaR and iFCPIs in *I. galbana* are labeled. **(E)** Superposition of the *V. stellata* PSI–VCPI and *T. minus* PSI–XLH structures. PsaS, PsaR and XLHs in *T. minus* are labeled. **(F)** Superposition of the *V. stellata* PSI–VCPI and *M. pyrifera* PSI–FCP structures. PsaR and FCPs in *M. pyrifera* are labeled.

**A**

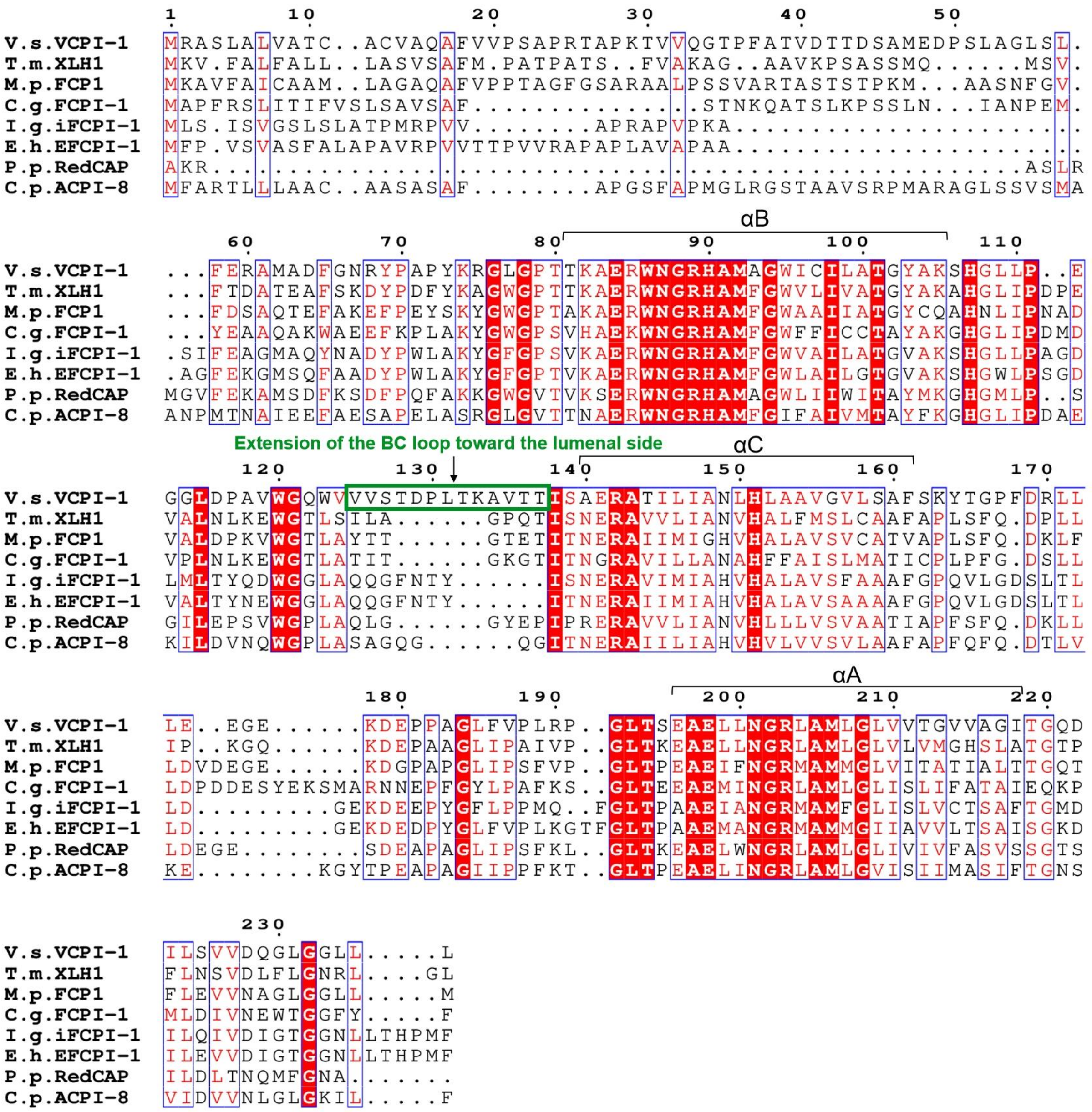

αB
αC
αA
Extension of the BC loop toward the lumenal side
V.s.VCPI-1 MRASLALLVATC..ACVAQAFVVPSAPRTAPKTVVQGTPFATVDTTDSAMEDPSLAGLSL.
T.m.XLH1 MKV.FALFALL..LASVSAFM.PATPATS..FVAKAG..AAVKPSASSMQ......MSV.
M.p.FCP1 MKAVFAICAAM..LAGAQAFVPPTAGFGSARAALPSSVARTASTSTPKM...AASNFGV.
C.g.FCPI-1 MAPFRSLITIFVSLSAVSAF.............STNKQATSLKPSSLN...IANPEM.
I.g.iFCPI-1 MLS.ISVGSLSLATPMRPVV.........APRAPVPKA.....................
E.h.EFCPI-1 MFP.VSVASFALAPAVRPVVTTPVVRAPAPLAVAPAA.....................
P.p.RedCAP AKR.......................................ASLR
C.p.ACPI-8 MFARTLLLAAC..AASASAF.......APGSFAPMGLRGSTAAVSRPMARAGLSSVSMA
V.s.VCPI-1 ...FERAMADFGNRYPAPYKRGLGPTTKAERWNGRHAMAGWICILATGYAKSHGLLP..E
T.m.XLH1 ...FTDATEAFSKDYPDFYKAGWGPTTKAERWNGRHAMFGWVLIVATGYAKAHGLIPDPE
M.p.FCP1 ...FDSAQTEFAKEFPEYSKYGWGPTAKAERWNGRHAMFGWAAIIATGYCQAHNLIPNAD
C.g.FCPI-1 ...YEAAQAKWAEEFKPLAKYGWGPSVHAEKWNGRHAMFGWFFICCTAYAKGHGLIPDMD
I.g.iFCPI-1 .SIFEAGMAQYNADYPWLAKYGFGPSVKAERWNGRHAMFGWVAILATGVAKSHGLLPAGD
E.h.EFCPI-1 .AGFEKGMSQFAADYPWLAKYGFGPTVKAERWNGRHAMFGWLAILGTGVAKSHGWLPSGD
P.p.RedCAP MGVFEKAMSDFKSDFPQFAKKGWGVTVKSERWNGRHAMAGWLIIWITAYMKGHGMLP..S
C.p.ACPI-8 ANPMTNAIEEFAESAPELASRGLGVTTNAERWNGRHAMFGIFAIVMTAYFKGHGLIPDAE
V.s.VCPI-1 GGLDPAVWGQWVVVSTDPLTKAVTTISAERATILIANLHLAAVGVLSAFSKYTGPFDRLL
T.m.XLH1 VALNLKEWGTLSILA......GPQTISNERAVVLIANVHALFMSLCAAFAPLSFQ.DPLL
M.p.FCP1 VALDPKVWGTLAYTT......GTETITNERAIIMIGHVHALAVSVCATVAPLSFQ.DKLF
C.g.FCPI-1 VPLNLKEWGTLATIT......GKGTITNGRAVILLANAHFFAISLMATICPLPFG.DSLL
I.g.iFCPI-1 LMLTYQDWGGLAQQGFNTY......ISNERAVIMIAHVHALAVSFAAAFGPQVLGDSLTL
E.h.EFCPI-1 VALTYNEWGGLAQQGFNTY......ITNERAIIMIAHVHALAVSAAAAFGPQVLGDSLTL
P.p.RedCAP GILEPSVWGPLAQLG......GYEPIPRERAVVLIANVHLLLVSVAATIAPFSFQ.DKLL
C.p.ACPI-8 KILDVNQWGPLASAGQG......QGITNERAIILIAHVHVLVVSVLAAFAPFQFQ.DTLV
V.s.VCPI-1 LE..EGE......KDEPPAGLFVPLRP..GLTSEAELLNGRLAMLGLVVTGVVAGITGQD
T.m.XLH1 IP..KGQ......KDEPAAGLIPAIVP..GLTKEAELLNGRLAMLGLVLVMGHSLATGTP
M.p.FCP1 LDVDEGE......KDGPAPGLIPSFVP..GLTPEAEIFNGRMAMMGLVITATIALTTGQT
C.g.FCPI-1 LDPDDESYEKSMARNNEPFGYLPAFKS..GLTEEAEMINGRLAMLGLISLIFATAIEQKP
I.g.iFCPI-1 LD.........GEKDEEPYGFLPPMQ..FGLTPAAEIANGRMAMFGLISLVCTSAFTGMD
E.h.EFCPI-1 LD.........GEKDEDPYGLFVPLKGTFGLTPAAEMANGRMAMMGIIAVVLTSAISGKD
P.p.RedCAP LDEGE........SDEAPAGLIPSFKL..GLTKEAELWNGRLAMLGLIVIVFASVSSGTS
C.p.ACPI-8 KE........KGYTPEAPAGIIPPFKT..GLTPEAELINGRLAMLGVISIIMASIFTGNS
V.s.VCPI-1 ILSVVDQGLGGLL.....L
T.m.XLH1 FLNSVDLFLGNRL.....GL
M.p.FCP1 FLEVVNAGLGGLL.....M
C.g.FCPI-1 MLDIVNEWTGGFY.....F
I.g.iFCPI-1 ILQIVDIGTGGNLLTHPMF
E.h.EFCPI-1 ILEVVDIGTGGNLLTHPMF
P.p.RedCAP ILDLTNQMFGNA.......
C.p.ACPI-8 VIDVVNLGLGKIL.....F

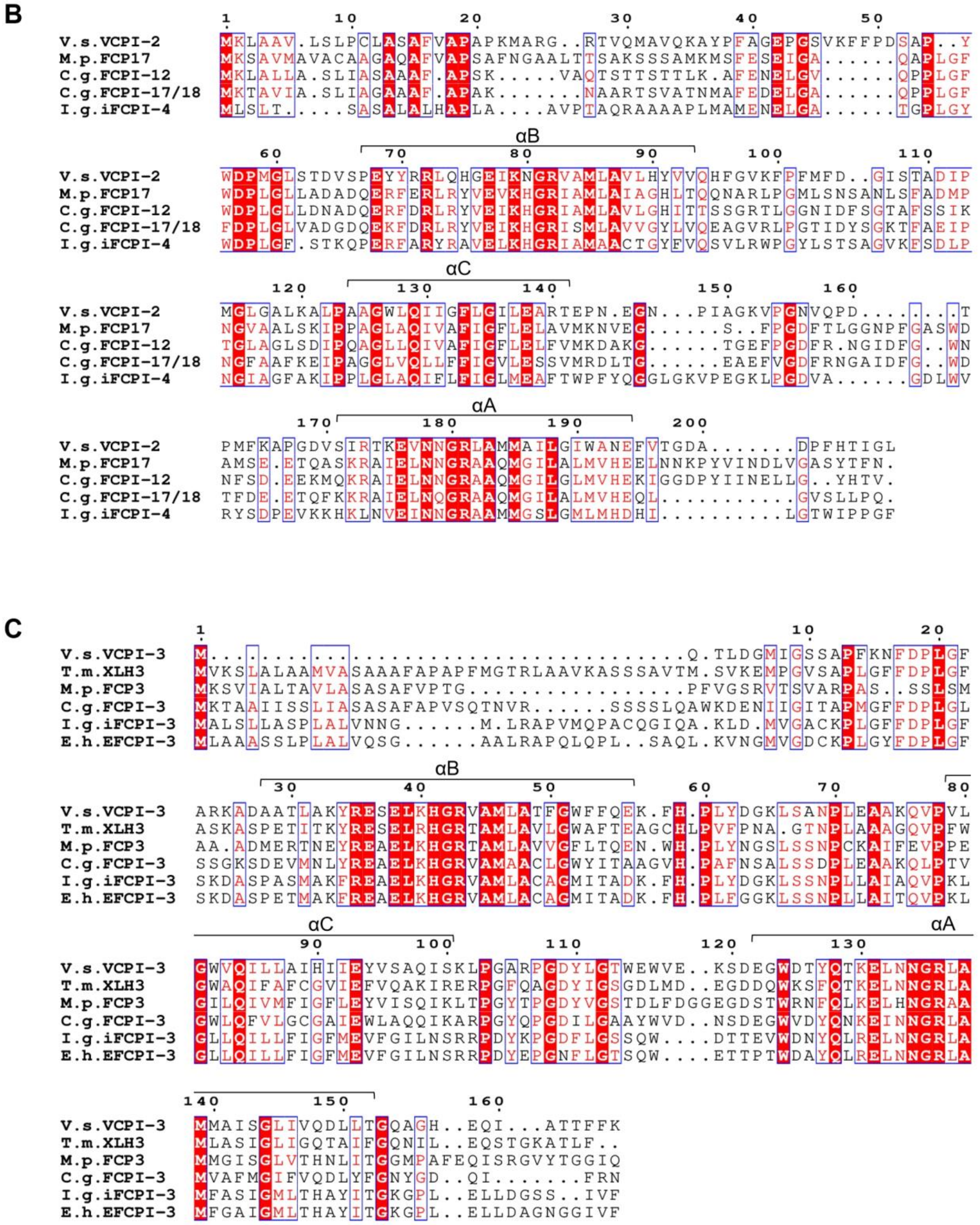

**Fig. S12. Sequence alignments of VCPIs with homologous LHCs from other red-lineage algae. (A)** Multiple sequence alignment of *Vischeria stellata* (*V.s.*) VCPI-1 with *Tribonema minus* (*T.m.*) XLH1, *Macrocystis pyrifera* (*M.p.*) FCP1, *Chaetoceros gracilis* (*C.g.*) FCPI-1, *Isochrysis galbana* (*I.g.*) iFCPI-1, *Emiliania huxleyi* (*E.h.*) EFCPI-1, *Porphyridium purpureum* (*P.p.*) RedCAP, and *Chroomonas placoidea* (*C.p.*) ACPI-8. **(B)** Multiple sequence alignment of *Vischeria stellata* (*V.s.*) VCPI-2 with *Macrocystis pyrifera* (*M.p.*) FCP17, *Chaetoceros gracilis* (*C.g.*) FCPI-12 and FCPI-17/18, and *Isochrysis galbana* (*I.g.*) iFCPI-4. **(C)** Multiple sequence alignment of *Vischeria stellata* (*V.s.*) VCPI-3 with *Tribonema minus* (*T.m.*) XLH3, *Macrocystis pyrifera* (*M.p.*) FCP3, *Chaetoceros gracilis* (*C.g.*) FCPI-3, *Isochrysis galbana* (*I.g.*) iFCPI-3, and *Emiliania huxleyi* (*E.h.*) EFCPI-3.

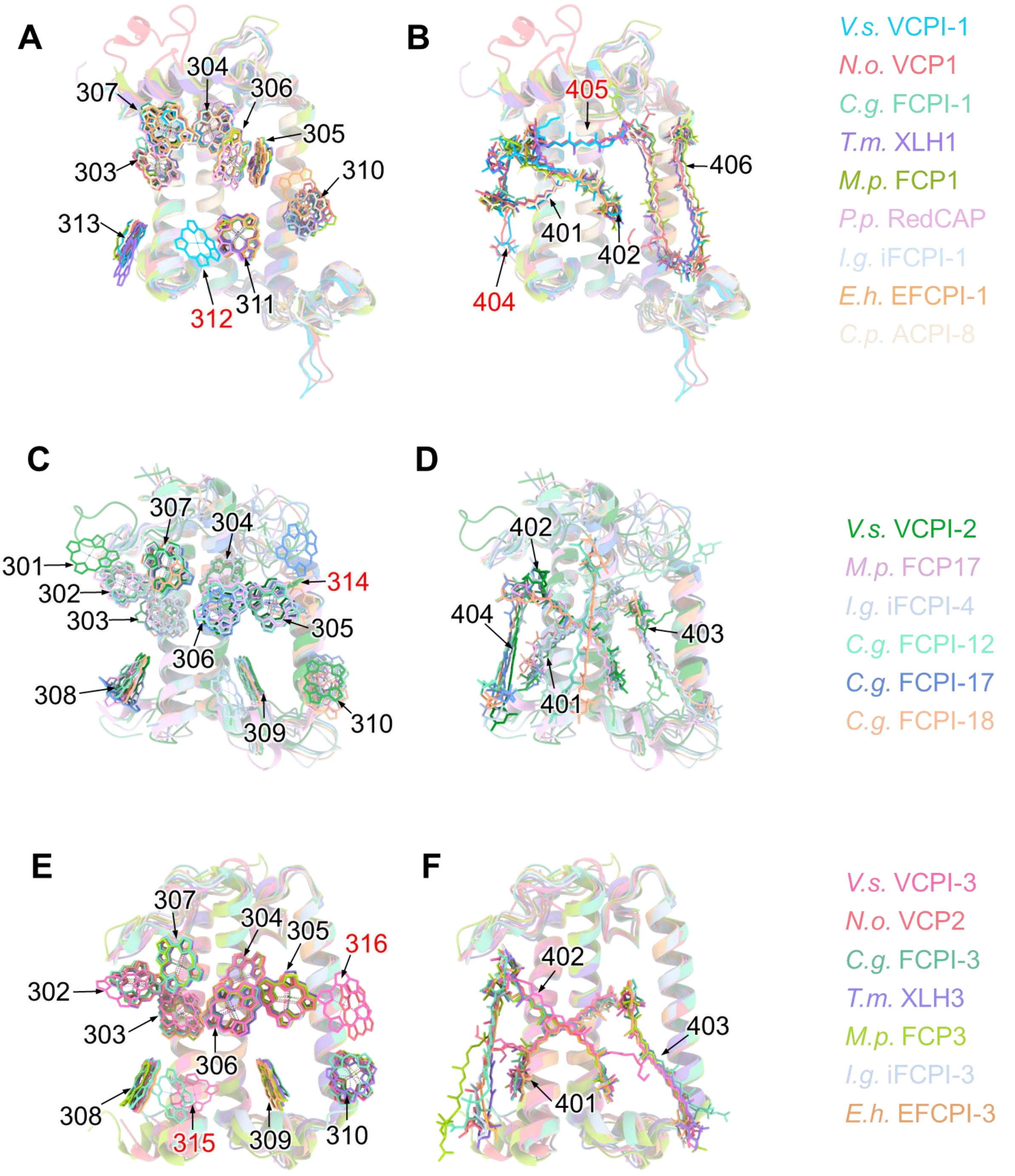

**Fig. S13. Comparison of pigment sites between VCPIs and LHCs from other red-lineage algae. (A to B)** Superposition of the Chl sites (A) and Car sites (B) in *Vischeria stellata* (*V.s.*) VCPI-1 with those in *Nannochloropsis oceanica* (*N.o.*) VCP1, *Chaetoceros gracilis* (*C.g.*) FCPI-1, *Tribonema minus* (*T.m.*) XLH1, *Macrocystis pyrifera* (*M.p.*) FCP1, *Porphyridium purpureum* (*P.p.*) RedCAP, *Isochrysis galbana* (*I.g.*) iFCPI-1, *Emiliania huxleyi* (*E.h.*) EFCPI-1, and *Chroomonas placoidea* (*C.p.*) ACPI-8. **(C to D)** Superposition of the Chl sites (C) and Car sites (D) in *Vischeria stellata* (*V.s.*) VCPI-2 with those in *Macrocystis pyrifera* (*M.p.*) FCP17, *Isochrysis galbana* (*I.g.*) iFCPI-4, and *Chaetoceros gracilis* (*C.g.*) FCPI-12, FCPI-17, and FCPI-18. **(E to F)** Superposition of the Chl sites (E) and Car sites (F) in *Vischeria stellata* (*V.s.*) VCPI-3 with those in *Nannochloropsis oceanica* (*N.o.*) VCP2, *Chaetoceros gracilis* (*C.g.*) FCPI-3, *Tribonema minus* (*T.m.*) XLH3, *Macrocystis pyrifera* (*M.p.*) FCP3, *Isochrysis galbana* (*I.g.*) iFCPI-3, and *Emiliania huxleyi* (*E.h.*) EFCPI-3. The compared antenna subunits are colored as indicated.

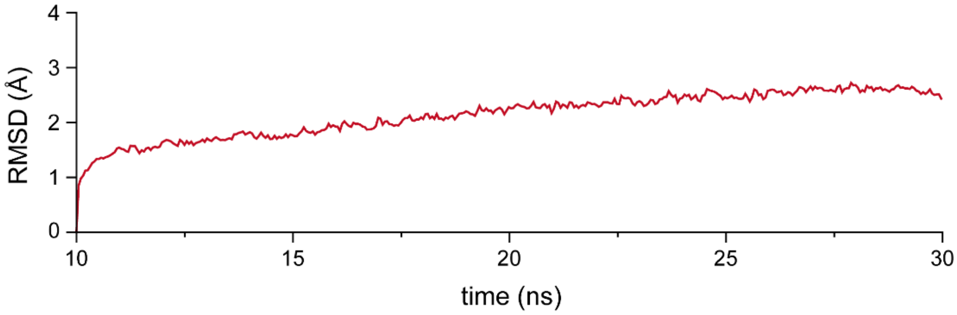

**Fig. S14. Root-mean-square deviation (RMSD) of the protein backbone for the protein-membrane system during the 10-30 ns period of the NPT molecular dynamics simulations.** The RMSD profiles indicate that the system reached structural stability during 20-30 ns. Therefore, representative conformations used for subsequent calculations were extracted from this 10 ns equilibrium interval at 2 ns intervals, yielding one frame every 2 ns.

**Table S1. Cryo-EM data collection, refinement, and validation statistics.**

| | PSI–VCPI (EMDB-82074; PDB-43QU) |
|---|---|
| **Data Collection and Processing** | |
| Voltage (kV) | 300 |
| Electron exposure (e- /Å2) | 60 |
| Defocus range (um) | -1.0~-1.8 |
| Pixel size (Å) | 0.53 |
| Symmetry imposed | C1 |
| Initial particle images (no.) | 1,290,982 |
| Final particle images (no.) | 151,622 |
| Map resolution (Å) | 2.44 |
| FSC threshold | 0.143 |
| **Refinement** | |
| Model composition | |
| Non-hydrogen atoms | 28,593 |
| Protein residues | 2672 |
| Ligands | 150 |
| *B* factors ($Å^2$) | |
| Protein | 12.10 |
| Ligand | 10.75 |
| R.m.s. deviations | |
| Bond lengths (Å) | 0.009 |
| Bond angles (º) | 1.118 |
| Validation | |
| MolProbity score | 1.55 |
| Clashscore | 4.18 |
| Rotamer outliers (%) | 0.18 |
| Ramachandran plot | |
| Favored (%) | 94.96 |
| Allowed (%) | 4.36 |
| Disallowed (%) | 0.68 |

**Table S2. Cofactors in each subunit of the eustigmatophyte VsPSI–VCPI supercomplex.**

| Subunits | Traced residues | Chl *a* | Car | Lipid | Others |
|---|---|---|---|---|---|
| PsaA | 738 (8-745) | 44 | 4 *β*-Car | 2 PG | 1 PQN, 1 SF4 |
| PsaB | 684 (2-288, 320-474, 495-736) | 33 | 5 *β*-Car | 1 DGDG | 1 PQN |
| PsaC | 80 (2-81) | | | | 2 SF4 |
| PsaD | 130 (4-133) | | | | |
| PsaE | 58 (4-61) | | | | |
| PsaF | 159 (27-185) | 1 | | | |
| PsaI | 34 (1-34) | 1 | 1 *β*-Car | | |
| PsaJ | 39 (10-48) | 1 | 2 *β*-Car, 1 Vio | | |
| PsaL | 171 (2-172) | 3 | 2 *β*-Car | | |
| PsaM | 30 (1-30) | | 1 *β*-Car | | |
| VCPI-1 | 181 (56-236) | 9 | 4 Vio, 1 Vau-ester | | |
| VCPI-2 | 170 (34-203) | 11 | 3 Vio, 1 Vau-ester | | |
| VCPI-3 | 155 (6-160) | 11 | 1 Vio, 2 Vau-ester | | |
| FNR | 43 (50-92) | | | | |
| PSI–VCPI | 2672 | 114 | 15 *β*-Car, 9 Vio, 4 Vau-ester | 2 PG, 1 DGDG | 3 SF4, 2 PQN |

DGDG, digalactosyldiacyl glycerol; PG, phosphatidyl glycerol; PQN, phylloquinone; SF4, sulphur–iron cluster.

**Table S3. Binding sites of pigments in the 3 LHC subunits of the eustigmatophyte VsPSI–VCPI supercomplex.**

| Sites | VCP-1 | VCP-2 | VCP-3 |
|---|---|---|---|
| 301 | | Chl *a* | |
| 302 | | Chl *a* | Chl *a* |
| 303 | Chl *a* | Chl *a* | Chl *a* |
| 304 | Chl *a* | Chl *a* | Chl *a* |
| 305 | Chl *a* | Chl *a* | Chl *a* |
| 306 | Chl *a* | Chl *a* | Chl *a* |
| 307 | Chl *a* | Chl *a* | Chl *a* |
| 308 | | Chl *a* | Chl *a* |
| 309 | | Chl *a* | Chl *a* |
| 310 | Chl *a* | Chl *a* | Chl *a* |
| 311 | Chl *a* | | |
| 312 | Chl *a* | | |
| 313 | Chl *a* | | |
| 314 | | Chl *a* | |
| 315 | | | Chl *a* |
| 316 | | | Chl *a* |
| 401 | Vio | Vio | Vau-ester |
| 402 | Vau-ester | Vio | Vau-ester |
| 403 | | Vau-ester | Vio |
| 404 | Vio | Vio | |
| 405 | Vio | | |
| 406 | Vio | | |

**Table S4. Structural information of the constructed protein-membrane simulation systems.** The dimensions of the simulation boxes, numbers of POPC molecules, water molecules, counterions, and total atoms for the three protein–membrane systems are summarized.

| | Dimension |
|---|---|
| Box dimensions, $Å^3$ | 172.39*169.20*127.62 |
| POPC lipids | 375 |
| Water molecules | 46914 |
| $Na^+$ ions | 150 |
| $Cl^-$ ions | 160 |
| Total atoms | 286841 |

**Table S5. Molecular dynamics equilibration protocol and positional restraints applied during system relaxation.** The two-stage equilibration procedure used for the protein-membrane systems is summarized. During lipid pre-equilibration, protein backbone atoms and retained pigments/cofactors were restrained to preserve the initial protein-cofactor architecture while allowing lipid, solvent, and ion relaxation. During whole-system equilibration, positional restraints were gradually released, followed by further NVT and NPT equilibration of the complete system. The restraint scheme corresponds to the molecular dynamics protocol described in the Methods section.

| Stage | Simulation step | Ensemble | Duration | restrained atoms |
|---|---|---|---|---|
| Lipid pre-equilibration | Energy minimization | | 50000 steps | Protein backbone, cofactors |
| Lipid pre-equilibration | Equilibration | NVT | 1ns | Protein backbone, cofactors |
| Lipid pre-equilibration | Equilibration | NPT | 10ns | Protein backbone, cofactors |
| Whole-system equilibration | Energy minimization | | 50000 steps | Pigments, IRON/SULFUR CLUSTER |
| Whole-system equilibration | Equilibration | NVT | 1ns | Pigments, IRON/SULFUR CLUSTER |
| Whole-system equilibration | Equilibration | NPT | 10ns | Pigments, IRON/SULFUR CLUSTER |
| Whole-system equilibration | Equilibration | NPT | 20ns | IRON/SULFUR CLUSTER |
